# ROBUST AND EFFICIENT ACTIVE GENETICS GENE CONVERSION IN THE RAT AND MOUSE

**DOI:** 10.1101/2022.08.30.505951

**Authors:** Chenyen Lai, Oscar Alvarez, Kristen Read, Don van Fossan, Christopher M. Conner, Shannon (Xaing-Ru) Xu, Dale O. Cowley, Valentino Gantz, David R. Webb, Kurt Jarnagin

## Abstract

The utility of Active Genetic (AG) gene conversion systems in rats and mice holds great promise for facilitating the production of complex strains harboring multiple humanizing genes. The practical application of such systems requires the identification of a robust, reusable, and highly efficient system. By characterizing twenty-eight different promoter and target site pairs we aimed to define the parameters needed to establish an efficient conversion system in male and female rats and mice. Using three specific meiosis prophase I active promoters to drive Cas9 expression. We studied several variables, including the number of Cas9 target sites, the distance between target sites, the cis versus trans configuration in linked pairs, and the effect of Cas9 copy number.

In the rat, three of twelve tested configurations provided efficient AG gene conversion in the 22% - 67% range, and four others catalyzed AG in the 0.7-1% range. The rat *Ddx4* (Vasa) promoter provides higher AG efficiency than the *Sycp1* promoter. In mice, ten of sixteen tested configurations, using the *Sycp1* and *pSycp1* promoters, provided efficiency in the 0.3% - 3.2% range. In rats, Cas9 expression levels are remarkably well correlated with AG gene conversion efficiency. The rat cis r*Cyp3A1*/r*Cyp3A2* locus was the most successful configuration, with gene conversion efficiencies of 0.7%-67%. This target site has a special property; the two Cas9 target sites are nearly perfectly homologous in the 100 bases around the gRNA target site.

Our findings identify key parameters that improve AG efficiency, including the use of two Cas9 target sites, and efficient promoters that drive high levels of Cas9 expression that are correctly timed during gamete development. These findings also uncover the unexpected benefit of high homology at paired gRNA target sites to promote efficiency. We provide new data to guide future efforts to develop yet further improved AG systems.

## Introduction

Multi-gene humanized^1^ rodent strains with human genes substituted for their rodent homologs have important applications for basic and applied research. Such strains can accelerate new drug discovery and development, promote understanding of the physiology arising from the interaction of several genes, and stimulate the development of animal disease models of autoimmunity, neurological diseases, and cancers. The creation of fully homozygous strains with more than four genes humanized is challenging due to the 1/2^n^ nature of Mendelian genetics. The fully homozygous product rate from a six-gene heterozygous-by-heterozygous cross is 1/4096, 0.02%; and for a ten-gene cross, the rate is 1/1,048,576. Using Mendelian genetics, creating multi-gene humanized mice or rats requires sequential steps stretching over years and the genotyping and handling of thousands of animals. Promotion of super-Mendelian inheritance by establishing a reusable, robust, and efficient^2^ Active Genetic (AG) gene conversion configuration^3^ in rats or mice offers a path to overcome these logistical obstacles by reducing the time and generations needed to create multiply humanized strains.

The foundation of Active Genetics in animals builds on genome editing approaches, first developed in *Drosophila*, applied in mosquitoes (Gantz and Bier, 2015; Gantz et al., 2015), and extended into mice (Grunwald et al., 2019). In this context, the Active Genetics configuration provides a Cas9^4^ source in combination with an AG cassette^5^ in gametes during developmental stages when Cas9 induced Double-strand breaks (DSBs) result in post-repair gene conversion. Due to the elevated rates of homology-directed repair (HDR) during meiosis (Ahmed et al., 2010; Xu and Greene, 2021), a chromosomal double-strand break (DSB) induced by Cas9 allows the AG cassette to be copied to the other wildtype chromosome, converting that element to homozygosity with super-Mendelian frequency. In a highly efficient system, the progeny should inherit the transgene 100% of the time instead of the Mendelian 50%. This form of Super-Mendelian inheritance would allow the efficient rapid assembly of large numbers of transgenes in a single individual. Once assembled, an array of efficient AG elements could be moved in a modular fashion into different genetic backgrounds within a few generations. With these strains, it would be possible to assemble complex combinations of genomic edits, including challenging closely linked alleles on the same chromosome. Using traditional approaches, creating higher order combinations becomes increasingly challenging, and humanizing linked genes is an inefficient, sequential, manpower, and time-intensive process.

Current rodent AG gene conversion systems achieved 5.5% efficiency^2^ in the female using a reusable Spo11 promoter and 42% efficiency^2^ using a single-use cre-lox-Vasa-driven Cas9 operating on the tyrosinase exon-4 locus and a mCherry AG cassette (Grunwald et al., 2019; Weitzel, et al., 2021). The Spo11 system also functioned with 3% efficiency in male mice (Weitzel, et al., 2021). These studies also suggested that the provision of Cas9-gRNA at or very near meiosis prophase I may be required to establish a working AG system (Grunwald et al., 2019; Grunwald et al., 2022). These studies employed a single trans-configured AG cassette, the tyrosinase cassette, using a cargo of 2.6 kb size and linked genetic marker 8.8 kb away to track gene conversion. This work inspired us to take the next step toward developing efficient rodent AG gene conversion systems and to better define the parameters necessary to make a practically useful system.

We begin by estimating the gene conversion efficiency required to prepare a practically useful and robust AG system. We aimed to define the parameters needed to prepare AG systems in both rats and mice, in males and females, and show its activity targeting loci and gene families of interest in cancer, immunology, and drug metabolism communities. We tested two target loci in two gene families, the *CYP450* family and the antibody-binding Fc-gamma receptor family (*FCGR*), including linked examples from each family. We also examined several variables, including the distance between AG sites, the configuration of AG sites – cis vs. trans, and the effect of the provision of homozygous versus heterozygous Cas9. Finally, we studied AG activity resulting from several different meiosis prophase-I active promoters to drive Cas9 expression. These studies validate the concept of active genetics in accelerating mammalian genetics and provide a framework for future studies to optimize these types of super-Mendelian inheritance systems.

## Methods

### Choice of loci for application of the AG process

One potential use of Active Genetic driven Super-Mendelian inheritance is to humanize large gene families in mammals such as rats or mice. Often gene families are genetically linked; thus, traditional Mendelian genetics makes humanization challenging due to the low recombination rates between linked genes. To demonstrate the application of Active Genetics in linked situations, we studied two gene families, the *hCYP450* family and the antibody-binding *hFc-gamma receptor* family (*hFCGR*). Due to their deep involvement in the development of antibody and small molecule therapeutics, both families are targets of multi-gene humanization by the drug metabolism and immunology communities (Scheer, et al., 2015; Petkova et al., 2006; Smith et al., 2012).

### Detection of Active Genetics Activity

To assess Active Genetics activity we crossed a strain bearing both Cas9 and a heterozygous Active Genetic cassette (AGC), the F1 parent, with a wild-type animal (Figure 1). Two methods were used to assess gene conversion activity: the Mendelian-excess test and the molecular marker test. The F2 offspring are evaluated via genomic analysis of tail DNA for the status of the locus on the receiver chromosome (Figure 1, lower) and the parentally derived chromosome. The Mendelian test uses the same assays as the molecular test described below. It quantifies AG statistically by detecting the presence of the AGC in excess over the Mendelian 50% expectation. The statistical surety of one’s conclusions is measured by a χ^2^ test using a p<0.05 figure of merit across about 200 offspring.

**Figure 1:**
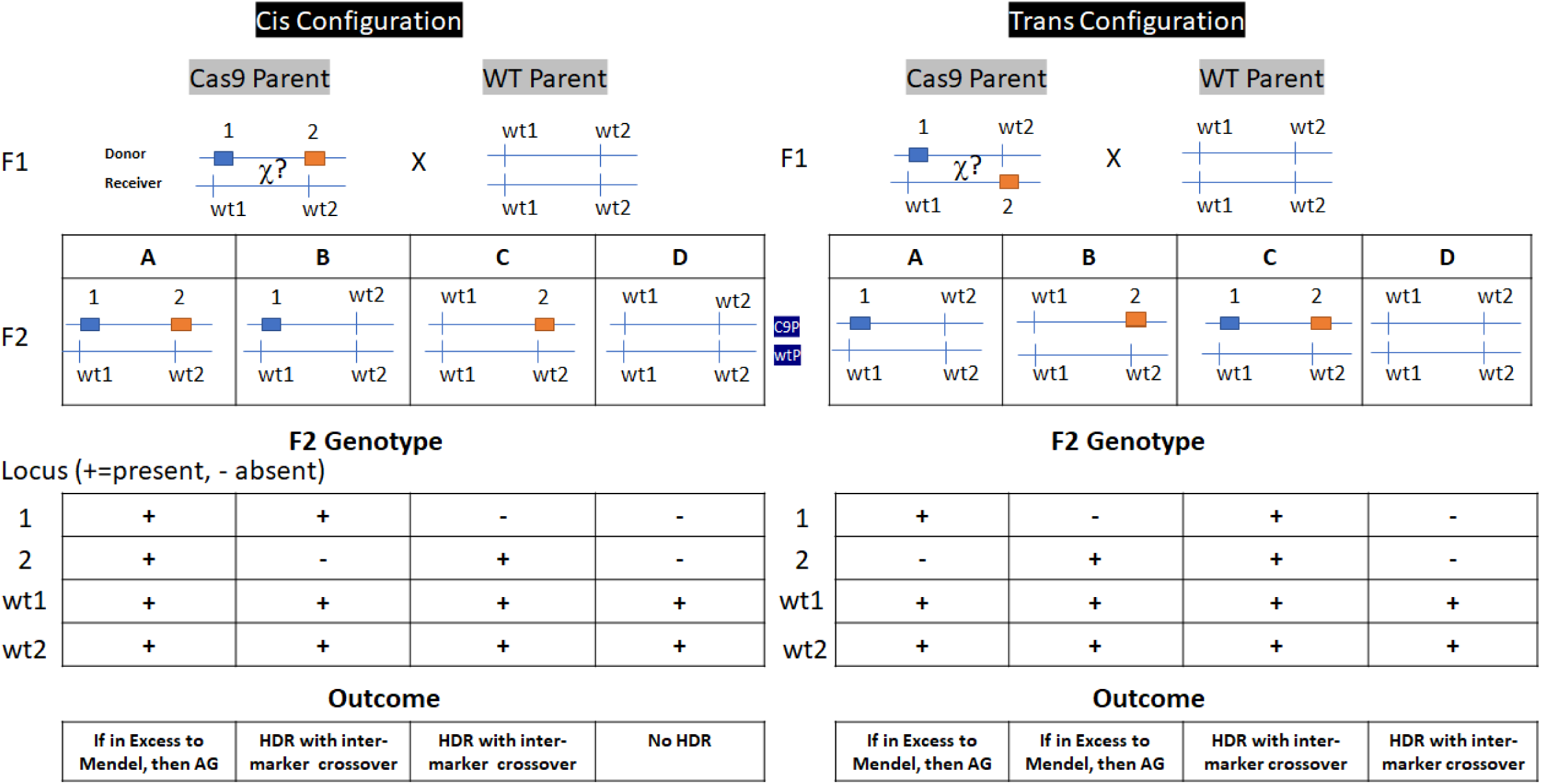
Test cross and outcomes guide for Cis or Trans configured AG gene conversion tests. AG gene conversion occurs in the gametes of the F1 Cas9 strain. Determination of the of F2 genotype assesses the outcome. The F1 cross is between a Cas9 plus AGC bearing F0 parent, C9P, and a wild type F0 parent, wtP. The four possible F2 outcomes are assessed by the presence or absence of the AGC or genetic marker, their frequency, and the status of the wild type loci. We measured AG gene conversion frequencies in two ways: the Mendelian Excess Test detects HDR terminated by crossover events outside the molecular markers; it assesses if there is an excess of AGC above the Mendelian expectation of 50%, condition A in the cis configuration, and condition A or B in the trans configuration. The molecular marker test assesses AG occurring via a DSB-induced repair event starting at one of the WT cut sites and subsequent termination of the HDR event by an inter-marker crossover. Molecular events are assessed by the observation of breakage of linkage condition B or C in the cis configuration, or development of linkage, condition C or D in the trans configuration. The method is applied to male or female F1 parents. On the donor chromosome, colored symbols, 1, or 2, are AGC or sequence markers, and wt1, and wt2 are the Cas9-gRNA target sites present on wild type-receiver-chromosome. χ?, represents a possible inter-marker crossover event that may happen during HDR-mediated repair of Cas9-caused DSBs. Not shown are HDR terminating crossover events outside the sequence markers, which may also occur.

In the molecular marker test (Figure 1), we prepared indels-markers or used naturally occurring genetic markers located from 1.7 kb to 83 kb from the humanized locus. Figure SM1 shows a typical molecular detection assay for the Rat *3A1-3A2* locus, and the supplemental materials provide further details on calculations and interpretation; the other cases were similarly arranged. Figure 1 provides a genotype interpretive guide for the genotypes observed during testing of cis or trans configured AGCs or sequence markers. In the molecular marker test, we assess genotypes by molecular biology and identify the HDR-caused transfer of the humanized locus to a receiver chromosome. We used a 2-fold redundant strategy for detecting molecular markers. First, hydrolysis probe-based quantitative PCR assays (TaqMan^®^) were conducted using probes specific for the humanized sequence at the gRNA target site on the humanized AGC bearing chromosome (donor chromosome) or the wild-type sequence at the gRNA target site (receiver chromosome). The probes used in these tests are specified in figure SM5. Second, we also scored a distantly located marker in the cis or trans configuration to the donor chromosome and scored the wild-type chromosome at that distant site. Thus, in net, we scored for the presence or absence of four alleles: the AGC, the wild-type sequence at the AGC site, the distant marker, and the wild-type sequence at the marker site. One can identify probable gene conversion events by measuring the presence or absence of these four sites and using Mendelian linkage reasoning based on the F2 cross conducted.

### Sequencing

We PCR amplified the wild-type locus at the gRNA target site (s), and Sanger sequenced the product in all F2 animals (primers in SM5). These sequencing reactions confirmed the hydrolysis probe findings and were analyzed using the ICE algorithm (Hsiau et al., 2018). ICE assesses the presence or absence of indels, the indel sequence, and the proportion of indel types seen on the receiver chromosome (lower chromosome in the figures). Indels are generated by Cas9 hydrolysis of the chromosome and subsequent non-specific end-joining (EJ) DNA repair mechanisms and attest to the presence of a biochemically active gRNA-Cas9 enzyme in the developing gamete, zygote, or embryo. EJ alleles represent the predominant competing DNA repair pathway countering the desired HDR outcome.

The frequency of indels, their sequence, and ratio to wild-type (WT) sequence provides an estimate of when during gametogenesis or embryo development at Cas9-gRNA action may have occurred. In the absence of an AGC, the presence of 50% WT and 50% of a single indel indicate activity during gametogenesis or at the single cell postfertilization stage. The presence of several distinct indels at approximately 25%, 12.5%, or 6.25% frequency suggests post-fertilization Cas9 activity at the four, eight, or more cell stages of embryo development, respectively (supplemental methods and results SR9A, SR9B.) From these findings, we can categorize Cas9-induced indels as occurring during gametogenesis, at the post-fertilization one or two-cell stage, as a post-fertilization event, or at the four or more-cell stage. These latter findings help us understand the success of our efforts to express biochemically active gRNA-Cas9 enzyme in the meiotic gamete and enzyme carryover or continued expression in the zygote or embryo.

### Strengths and Weaknesses of Gene-Conversion tests

These two gene conversion tests have distinct strengths and weaknesses. The Mendelian test is conceptually simple and can detect HDR events outside the molecular marker, which is a probable event due to linkage and crossover interference. The molecular test is unequivocal. Intermarker crossover and gene conversion happened if the humanized gene and the marker occur on the same chromosome in a trans configuration. In the cis configuration, dissociation of the markers indicates intermarker crossover. However, the molecular test is a low probability event because crossover during HDR-mediated gene conversion is not likely in small genetic regions of less than one centimorgan, about 1 million base pairs; this is the case for all the studied chromosomal markers (Figure 2), and the marker used in previous rodent studies (Weitzel et al., 2021; Pfitzner et al., 2020). The Mendelian excess test relies on statistical power assessed using a χ^2^ test. Our statistical power versus efficiency analysis (Supplemental methods) indicates that we need to evaluate about 200 offspring to achieve statistical significance in a setting where the Active Genetic efficiency is 25%. The number of offspring required to reach a statistically significant outcome versus decreasing efficiency is a steeply rising function; hence the Mendelian test has a practical lower bound in terms of sensitivity. Thus, the molecular test is an unequivocal and precise demonstration of a gene conversion event, but it could underestimate the actual gene conversion efficiency and thus is less sensitive than the genotype-based Mendelian test. The Mendelian test is a more accurate estimate of the overall gene conversion efficiency, but it has a lower bound on its detection power when AG gene conversion efficiency is low.

**Figure 2:**
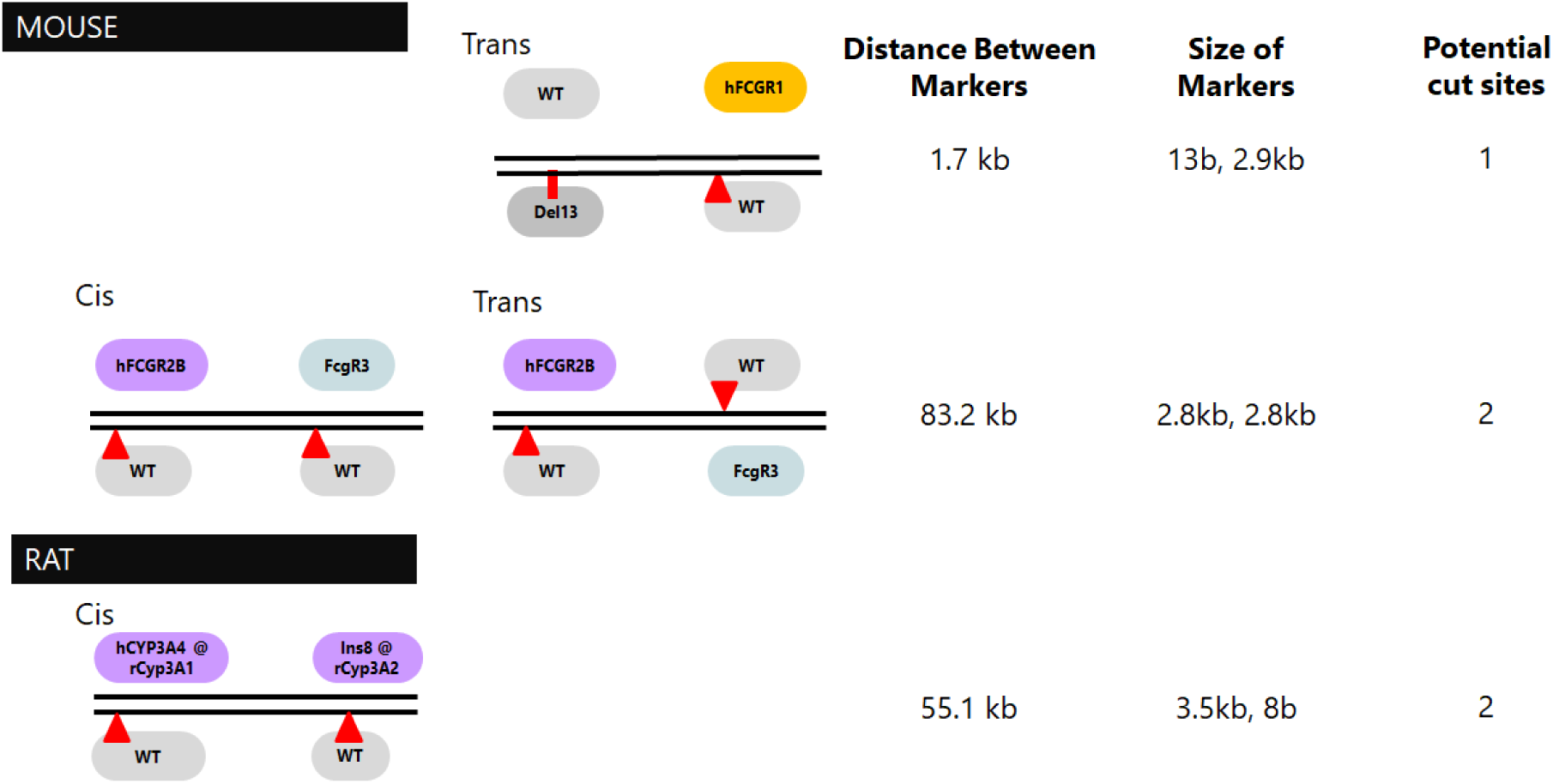
Arrangement of the Active Genetic Cassettes, target loci, gRNA cut sites, and chromosomal markers to allow Mendelian and molecular detection of Active Genetic events. Not shown is one additional construct, humanized rat 2E1, arranged with one N-terminal gRNA cut site and no marker on the chromosome; in this case, we detect Active Genetics by the Medelian excess method. The receiver chromosome is at the tip of the red triangle, which denotes the gRNA cut site. Red rectangle: indel maker, without a gRNA cut site. The humanized locus, non grey-colored symbols, contains an Active Genetic Cassette.

### Cas9 Absence Test

To ascertain the background rate of AG gene conversion events, we assessed the Mendelian frequency, or molecularly demonstrated events, in mice and rat strains with the humanized AG cassette present but no Cas9 gene. The Cas9 absence test was conducted for all four test loci, in the cis and trans configuration, and in male and female mice or rats.

### F1 strain identity and quality check

Genotypes of all F1 animals in Active Genetics tests were checked by DNA sequencing presence of the wildtype gRNA sites on the receiver chromosome and to show that the genotype planned was present. The test was also conducted to assure that during the creation of these strains, Cas9-induced indels did not occur. Indel-bearing parents were removed from testing; they are not capable of hosting AG events.

### Site-dependent configuration of Active Genetic cassettes

Previous work in mice examined a single configuration of a 2.6 kb Active genetic cassette at the tyrosinase exon-four locus and used a trans-3’-SNP marker located 8.8 kb away to detect gene conversion (Grunwald, et al., 2019; Weitzel et al., 2021; Pfitzner et al., 2020). Since a robust Active Genetic system is an important goal, we explored several loci. These loci differed in their chromosomal environment and distance to other active genetic target elements or indel markers. We also examined differences in performance when the AG cassettes were configured in chromosomal locations cis or trans to each other. Figure 2 shows the arrangement of loci examined.

### Configuration of humanized target loci, the Active Genetic Cassette

Humanization of the target loci followed a standardized model. Each rodent target locus was reconfigured to express the human cDNA for the target in exon 1 of the corresponding rodent gene. Efforts were made to insert the human initiation methionine and cDNA at, or close to, and in-frame with, the rodent initiation methionine. Once formed, these constructs insert an in-frame human minigene in exon 1 of the rodent gene and disrupt the rodent gene. Each AG humanization cassette also contains a guide RNA expression cassette with a U6 promoter element driving expression of the same guide RNA used to insert the human minigene into the mouse genome, thus targeting the same site on the wild-type mouse chromosome for AG gene conversion when combined with a Cas9 transgene. We prepared five AGC and genomic marker configurations (Figure 2) These include rats humanized with *hCYP2E1* with no genomic marker. They also include a Rat *Cyp3A1* humanized with *hCYP3A4* and a gRNA that targets both the *rCyp3A1* and *rCyp3A2* locus. This arrangement is possible because of the very high homology of these loci; they have perfect identity in the 20 bases on either side of the target site. Mouse humanized with *hFCGR2b* and *hFCGR3*a in the cis and trans configuration were also prepared. And finally, mouse *FcgR1* with a trans-Del13 genomic marker, or the identically located Del24 marker indel^6^. Due to the details of each gene’s sequence, each targeted locus includes slight deviations from these general plans as detailed in the supplemental methods (Figure SM4).

### Identification of candidate gene promoters

The search for potential promoter sequences to drive Cas9 expression in amounts and timed to promote Active Genetics was based on extensive literature investigations described in detail in the Supplemental Methods. Briefly, earlier mouse studies of meiosis-restricted promoters have used the Vasa (*Ddx4*), Stra8, and Spo11 gene promoters or studied synthetic non-meiosis-restricted constitutive promoters (Grunwald-Cooper, 2019; Weitzel-Cooper, 2020; Pfitzner-Thomas, 2021). Gene conversion efficiencies in these studies ranged from 0%-42% when applied to a test locus, tyrosinase exon-4, using single-use or reusable systems. The key finding indicates that Active Genetics in rodents is favored when Cas9 is expressed at or near meiosis prophase I. In our analysis we sought genes with high LZ/2C expression ratios (ratio of expression in leptotene-zygotene cells to 2x chromosomal DNA content cells) and lower expression levels in the 2C stage. We also focused on genes with prophase dominant expression, known involvement with chromosomal pairing or synaptonemal complexes, and recombination (Bolcun-Filas and Handel, 2018). We identified the candidate genes *Sycp1, Sycp2, Sycp3, Syce1, Rad51, Dmc1, Msh4, Msh5, Ddx4(Vasa/Mvh)*, and *Hfm1* that have LZ/2C expression ratios from 3 to 43. These genes, proteins, and promoters were further parsed based on literature findings as to their known restriction to spermatogonia and oogonial cells and the degree to which the promoter had been characterized for embryonic lethality and sterility in males and females. This analysis leads to the identification of the full length *Sycp1* promoter and its 824 bp promoter fragment, p*Sycp1* (Sage et al., 1999). The *Vasa* promoter (*Ddx4* in the rat and Vasa in humans, and also called *Mvh* in the mouse) was also selected because this promoter functioned to promote efficient Active Genetics in *Drosophila melanogaster* (Gantz, and Bier, 2015) and mosquitoes, *Anophiles gambiae* (Gantz et al., 2015). It also functioned in a single-use setting in mice, *Mus musculus* (Grunwald et al., 2019). This gene has an LZ/2C ratio of 13 and is restricted in expression to gametogonia cells during meiosis. (DaCruz et al., 2016; Toyooka, 2000).

### Preparation of Accelerator strains

Accelerator Strains were configured to express Cas9 at or near the time of meiosis, using a meiosis-restricted promotor or a drug-regulated Cas9 enzyme^7^. The supplemental methods and Figure SM1 provide construction detail and show the configuration of four rats or mouse Cas9 accelerator strains prepared. In general, strains were constructed with the mammalian expression codon-optimized sp*Cas9* gene integrated behind the target promoter and its protein cDNA. A polyprotein was formed encoding the target protein, often separated by the P2A self-cleavage sequence before the sp*Cas*9 gene, rat *Ddx4*, and rat and mouse *Sycp1* genes. A co-cistronic green-fluorescent protein was also provided downstream from Cas9 and separated by a peptide self-cleavage sequence to assist tissue expression studies. In other cases, the *Sycp1* promoter fragment, p*Sycp1*, was placed upstream of sp*Cas*9 and was integrated into intron-1 of the *Rosa*26 gene in mice.

## Results

### Efficiency

As a starting point for our studies, we addressed the question: What efficiency of formation of fully homozygous offspring is needed to provide a practically useful method? We examined a heterozygous by heterozygous cross, Aa X Aa. We define the Mendelian frequency of fully homozygous offspring as a percentage (M%) as M%=100 × 1/2^n^ × 1/2^n^, where n is the number of loci. We define Active Genetic efficiency, AG%, as 100 times the frequency that the parental genotype is copied to the receiver chromosome (Table 1). When driven by Active Genetics, the overall percent efficiency at which all loci become homozygous in one breeding cycle is defined as Hom%, and it is calculated by: Hom% = (100 × (AG%/100)^n^) + (M% × (1-AG%/100)). Table 1 shows this calculation carried out over AG gene conversion efficiency ranging from 100% to 25%, addressing 1 to 10 loci under the simplifying assumption of equal and independent conversion efficiencies at all loci, and we assumed that male and female parents are equally AG capable. The Mendelian homozygosity yield drops precipitously as the number of loci increases. Similarly, the homozygous yield decreases sharply as the AG efficiency decreases (Table 1). From this analysis we conclude that the Active Genetic process needs to reach an efficiency greater than 70%, to be practically useful for the assembly of more than 5 loci. Furthermore, the process must be robust, highly efficient at all loci, and reusable in successive generations.

**Table 1:**
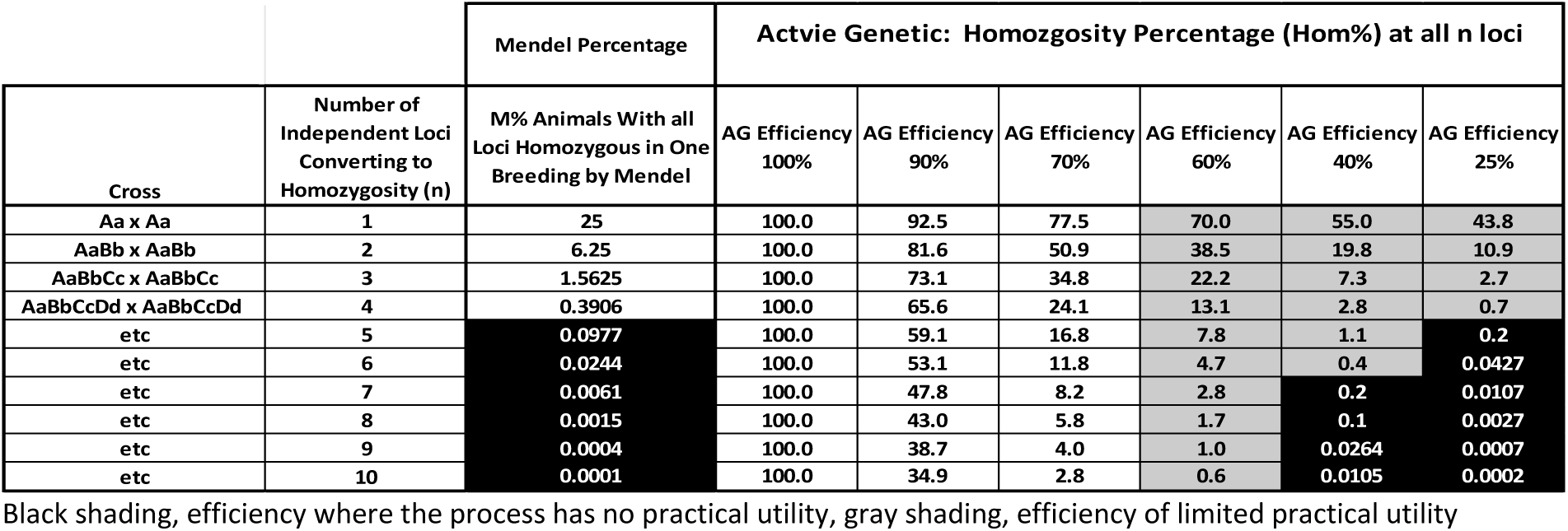
Percentage of animals full homozygous in a heterozygous-by-heterozygous cross versus Active Genetic efficiency, and versus the number of heterozygous loci.

| Cross | Number of Independent Loci Converting to Homozygosity (n) | Mendel Percentage | Active Genetic: Homozygosity Percentage (Hom%) at all n loci |  |  |  |  |  |
| --- | --- | --- | --- | --- | --- | --- | --- | --- |
|  |  | M% Animals With all Loci Homozygous in One Breeding by Mendel | AG Efficiency 100% | AG Efficiency 90% | AG Efficiency 70% | AG Efficiency 60% | AG Efficiency 40% | AG Efficiency 25% |
| Aa x Aa | 1 | 25 | 100.0 | 92.5 | 77.5 | 70.0 | 55.0 | 43.8 |
| AaBb x AaBb | 2 | 6.25 | 100.0 | 81.6 | 50.9 | 38.5 | 19.8 | 10.9 |
| AaBbCc x AaBbCc | 3 | 1.5625 | 100.0 | 73.1 | 34.8 | 22.2 | 7.3 | 2.7 |
| AaBbCcDd x AaBbCcDd | 4 | 0.3906 | 100.0 | 65.6 | 24.1 | 13.1 | 2.8 | 0.7 |
| etc | 5 | 0.0977 | 100.0 | 59.1 | 16.8 | 7.8 | 1.1 | 0.2 |
| etc | 6 | 0.0244 | 100.0 | 53.1 | 11.8 | 4.7 | 0.4 | 0.0427 |
| etc | 7 | 0.0061 | 100.0 | 47.8 | 8.2 | 2.8 | 0.2 | 0.0107 |
| etc | 8 | 0.0015 | 100.0 | 43.0 | 5.8 | 1.7 | 0.1 | 0.0027 |
| etc | 9 | 0.0004 | 100.0 | 38.7 | 4.0 | 1.0 | 0.0264 | 0.0007 |
| etc | 10 | 0.0001 | 100.0 | 34.9 | 2.8 | 0.6 | 0.0105 | 0.0002 |

Black shading, efficiency where the process has no practical utility, gray shading, efficiency of limited practical utility

### Cas9 Expression

#### mRNA analysis of Cas9 expression in Accelerator Strains

We examined Cas9 mRNA expression in the organs of rat and mouse Accelerator Strains in post-puberty testis and ovary and a non-gonadal tissue, the kidney (Figure SR1). In nongonadal tissue, the kidney, we observed a very low level of Cas9 mRNA, except in the case of the mouse strain with a partial promoter p*Sycp1*, where expression occurred in gonadal and non-gonadal tissues. There was less mRNA expression detected in the post-puberty ovary of all strains. Cas9 mRNA was strongly expressed in the testis of rats and mice with all four promoters, Figure SR1. In summary, our findings reveal that Cas9 mRNA is expressed at lower levels in adult ovaries and much higher levels in the testis of Accelerator rats and mice.

### Immunohistochemical detection of Cas9 in Accelerator Strains

Rat accelerator strains with Cas9 regulated by the *Ddx4* (Vasa) promoter and the *Sycp1* promoter were examined for expression of Cas9 in the testis and ovary. Table 2 summarizes those findings derived from figures SR2-5. Figure SR2 shows that Cas9 was strongly expressed in the rat testis when driven by the *Ddx4* or *Sycp1* promoter. Both promoters drove strong expression in the cells of the size, position, and shape corresponding to pachytene primary spermatocytes through post-meiotic II secondary spermatocyte stage of spermatogenesis. We observed prominent accumulation in the chromatid body, especially in the *Ddx4*-driven animals. Cas9 was present in both the cytoplasm and nucleus. In either strain, expression was weaker in the spermatogonial cell layer but slightly more prominent in the *Ddx4*-driven strain. The expression patterns for rat *Sycp1* and *Ddx4*-driven expression were similar to our design targets.

**Table 2:** Immunohistochemical assessment of the expression of Cas9 in Testis of Accelerator Strains, images figure SR2-5.

| Species<br>Promoter | Stain<br>Intensity<br>Het | Stain<br>Intensity<br>Hom | Cell layer with<br>Prominent<br>expression | Location of intracellular<br>expression |
| --- | --- | --- | --- | --- |
| Mouse<br>*pSycp1* | +++ | +++ | Round spermatids<br>and potentially 2°<br>spermatocyte,<br>elongating<br>spermatids | Prominent cytoplasm,<br>slight nucleus |
| Mouse-<br>*Sycp1* | ++ | +++ | Spermatogonia | Prominent nuclear, slight<br>cytoplasm |
| Rat<br>*Sycp1* | ++ | +++ | Spermatogonia | Nucleus and cytoplasm;<br>Slightly more prominent<br>in chromatid bodies |
| Rat<br>*Ddx4* | ++ | N/A | 1° spermatocytes<br>through round<br>spermatids | Nucleus and cytoplasm;<br>Prominent in chromatid<br>bodies |

Het, heterozygous; Hom, homozygous; grey shading rat; white shading mouse

In mice, both the Sycp1 and pSycp1 promoters drive significant Cas9 expression but in different cell types (Figure SR3). The pSycp1 promoter mainly induced expression in post-meiosis I cells, exhibiting the size, position, and shape of secondary spermatocytes and spermatids. However, some cells of the size, shape, and location of pachytene cells may also have been stained (SR3-panel C). The Sycp1 promoter strain expression of Cas9 was sharply confined to the spermatogonia cell layer in the cytoplasm and nucleus (SR3 panels D and E).

*Sycp1-Cas9*-driven expression in rats and mice was highly similar; in both species, the promoter and Cas9 configuration were almost identical (SR2-B and C vs. SR3-D and E). In the *Sycp1-Cas9* mouse and rat strains, the *Sycp1*-driven Cas9 expression was most prominent in cells of the spermatogonia layer with slight carryover into primary spermatocytes. In the *Sycp1*-Cas9 rat and mouse strains, Cas9 was present in cells similar to primary spermatocytes base on the shape, size, and position of the cells. Cas9 expression extended through post-meiotic II round spermatids. There is prominent accumulation in the chromatid body, especially in the rat.

### Homozygous configuration

(Figure SR5) shows additional Cas9 expression for the homozygous rat and mouse *Sycp1* strains. When homozygous, the *Ddx4*-Cas9 rat strain was infertile, azoospermic and exhibited staining of the spermatogonia similar to WT control staining. The mouse pSycp1-Cas9 transgene expression was not appreciably different in heterozygotes versus homozygotes.

### Cas9 in the developing sperm

The cell stage during meiosis I at which Cas9 expression occurs was further characterized by assessing the DNA content of testicular cells and Cas9 content using flow cytometry. This method allowed us to distinguish 1N, 2N, and 4N DNA content cells and simultaneously the content of the surrogate marker, Cas9-T2A-GFP (Figures SR6A-B and Figures SR6C-E). The use of GFP as a surrogate marker comes with a caveat. Our Cas9 constructs have a C-terminal NLS, whereas our GFP protein does not; hence the fates of NLS-tagged-Cas9 and GFP may not overlay precisely with the immune-histochemical detection of NLS-tagged-Cas9. At the cellular level, they provided broadly similar findings, as shown by the immune-histochemistry of serial sections in Figure SR7

Immunohistochemical localization of Cas9 in the male testis of *Ddx4-Cas9* rats provides clear evidence of expression in the target cell population, from 1° spermatocytes through round spermatid stages. Cas9 mRNA is also present in female ovaries of these rat strains, and immunohistochemical detectable Cas9 is still present in nearly mature eggs of *Ddx4* and *Sycp1* rats. Mouse *Sycp1*-Cas9-T2A-GFP animals express Cas9 and the GFP surrogate marker similarly to the rat *Sycp1*, including immuno-histochemical and GFP surrogate marker detection in 1N, 2N, and 4N DNA content cells. Mouse p*Sycp1*-Cas9-T2A-GFP expression occurs in a more diffuse pattern than in rats or mice with other configurations. The highest levels detected were in post-meiotic cells, round spermatids, and spermatocytes. However, there are also GFP bright cells with 4N DNA content. Rats or mice homozygous for Cas9 encoding constructs express higher levels of Cas9 protein than heterozygous animals. In *rSycp1-*P2A-GFP rats, 1N, 2N, and 4N DNA content cells exhibited substantially greater levels of GFP expression than wild-type controls, and all the DNA populations displayed many more GFP bright cells than wild-type animals. Homozygous *rSycp1*-T2A-GFP animals expressed higher levels of GFP than heterozygous animals (SR6A). Using direct Cas9 staining, rDdx4-Cas9 animals showed more Cas9 staining than wild-type animals, and the 4N populations contain 6-fold more Cas9 positive cells than wild-type animals (SR6B). In mice, both m*Sycp1-*T2A-Cas9 and m*pSycp1-* Cas9 -P2A-GFP animals showed strong expression of GFP in the total testicular cell population compared to wild-type animals (SR6A). There were more GFP-positive cells in the 4N cell population than in the 4N wild-type testis. Homozygous animals had larger 4N GFP positive populations than heterozygous animals. In mice, the 1N and 2N cell populations showed similar patterns. Our immunocytochemical detection revealed that 4N cells in leptotene through pachytene expressed Cas9-GFP.

In summary, the mRNA, immunohistochemical, and flow cytometric studies of Cas9 provide an overall view that expression of Cas9 is occurring and at substantial levels in developing sperm. Expression in 4N cells during the leptotene through the pachytene stage was observed. Expression at this stage is a key cell stage targeted by our constructs. Other developmental stages also express Cas9, including pre-meiosis and post-meiosis stages. In all the strains, immunocytochemical and immunohistochemical staining displayed higher levels of Cas9 in homozygous animals and less but detectable levels in heterozygous animals.

### Biochemical evidence of Cas9 activity in gonadal cells

To detect biochemically active Cas9 in the testis of rats, we performed a T7 endonuclease 1 (T7E1) mismatch detection assay following the details provided in the supplemental methods (Li, et al., 2013; Marshal et al., 1995; reviewed in Sentmanat et al., 2018). In both the rat *Ddx4*-Cas9 and *Sycp1*-Cas9 strains containing an AGC encoded gRNA, but not when Cas9 was absent, testicular DNA contains end-joining (EJ) repaired indels at the location of the gRNA target site (Figure SR8). This result confirms the presence of biochemically active Cas9 in the rat testis.

### Biochemical activity as shown by indels

Examining the presence or absence of indels at the gRNA target site in F2 AG test animals provides evidence for the presence of gRNA-Cas9 and its biochemical activity in the gamete or zygote (Table SR9A, SR99B). Indels, which form as a result of the EJ repair mechanism following gRNA-Cas9 hydrolysis of the chromosome. We observed rates in the F2 offspring ranging from 0.5%-51% in rats (Table SR9A) and 0.8%-47% in mice (Table SR9B). Gene conversion and indel events resulted from gRNA-Cas9-induced DSBs since no indels and no AG gene conversion events were observed in strains encoding the AG cassette with its gRNA but without any Cas9 gene (Table SR9A and SR9B). Interestingly, mouse indel formation rates were significantly higher in males than females. Offspring of male mice averaged 29.7% ± 6.9% (N=8) indels while offspring of female mice averaged 7.2% ± 7.3 (N=7) indels in F2 pups, p<0.05. No similar male-to-female skew was noted in rats. The difference in Cas9 efficiency in male and female mice suggests that there may be subtle changes in the activity of Cas9 and efficiencies of the EJ pathways in developing gametes of each sex and species. Additional studies will be needed to understand these phenomena. The AGC-bearing rat and mouse strains examined carried indels indicating that biochemically active gRNA-Cas9 was present in developing gametes, zygotes, or embryos (SR9A and SR9B). For example, the heterozygous female *Ddx4* accelerator strain had 30 of 526 pups, and heterozygous males had 31 of 224 pups, wherein the AGC was absent, and 50% of a single indel and 50% of a WT sequence were present. This pattern is most consistent with Cas9 acting in the gamete or one-cell post-fertilization stage.

### Active Genetics gene conversion

Three of the twelve tested configurations in the rat provided efficient AG gene conversion in the 22-67% range, and four others catalyzed less robust AG in the 0.7%-1% range, as demonstrated using the molecular test (Table 3A). None of the sixteen mouse strains provided highly efficient AG gene conversion; however, ten of the sixteen strains provided less robust AG with efficiency in the 0.8 – 3.2% range, as demonstrated using the molecular test (Table 3B, 3C).

**Table 3A:** Rat AG gene conversion efficiency: the influence of Cas9 promoter, Cas9 gene dosage, locus, and parent sex.

| Rat (8/2) |  |  |  |  | AG Gene Conversion |  |  |  |  |  |  |  |
| --- | --- | --- | --- | --- | --- | --- | --- | --- | --- | --- | --- | --- |
|  | A | B | C | D | E | F | G | H | I | J | K | K |
|  | Promoter-Cas9 | Cas9 Hetero or Homozygous | AG cassette gRNA target site (number of sites) | Donor chromosome Genetic Marker | Sex | Number Pups Examined | Intermarker Molecular Biology Events observed (B,C) | Number of Observed AGC Present, Including Molecu (A+B+C) | % Gene conversion (AG Efficiency), Including Molecular | x2 p-val | % Gene Conversion Significance (n.s., p>0.05) | Interperated Amount of AG Observed? |
| 1 | Ddx4-Cas9 | Hetero | rCYP2E1 (1) | hCYP2E1 | M | 207 | n/a | 107 | 3.4% | 0.73 | n.s. | n.s. |
| 2 |  | Hetero | rCYP2E1 (1) | hCYP2E1 | F | 186 | n/a | 102 | 9.7% | 0.35 | n.s. | n.s. |
| 3 |  | Homo | rCYP2E1 (1) | hCYP2E1 | F | 125 | n/a | 75 | 20.0% | 0.11 | n.s. | n.s. |
| 4 |  | Hetero | rCYP3A1, r3A2 (2) | cis-hCYP3A4, Ins8 | M | 224 | 2, 2 | 121 | 8.0% | 0.40 | n.s. | 0.9% |
| 5 |  | Hetero | rCYP3A1, r3A2 (2) | cis-hCYP3A4, Ins8 | F | 526 | 10, 11 | 408 | 55.1% | 3.9E-19 | 3.9E-19 | 55.1% |
| 6 |  | Homo | rCYP3A1, r3A2 (2) | cis-hCYP3A4, Ins8 | F | 205 | 3, 10 | 171 | 66.8% | 1.3E-11 | 1.3E-11 | 66.8% |
| 7 | Sycp1-P2A-Cas9 | Hetero | rCYP2E1 (1) | hCYP2E1 | M | 211 | n/a | 100 | -5.2% | 0.59 | n.s. | n.s. |
| 8 |  | Hetero | rCYP2E1 (1) | hCYP2E1 | F | 211 | n/a | 114 | 8.1% | 0.41 | n.s. | n.s. |
| 9 |  | Hetero | rCYP3A1, r3A2 (2) | cis-hCYP3A4, Ins8 | M | 206 | 2, 2 | 104 | 1.0% | 0.92 | n.s. | 1.0% |
| 10 |  | Homo | rCYP3A1, r3A2 (2) | cis-hCYP3A4, Ins8 | M | 336 | 4, 2 | 186 | 10.7% | 0.16 | n.s. | 0.9% |
| 11 |  | Hetero | rCYP3A1, r3A2 (2) | cis-hCYP3A4, Ins8 | F | 210 | 0, 3 | 102 | -2.9% | 0.77 | n.s. | 0.7% |
| 12 |  | Homo | rCYP3A1, r3A2 (2) | cis-hCYP3A4, Ins8 | F | 164 | 3, 5 | 100 | 22.0% | 0.047 | 0.047 | 22.0% |
| 13 |  | NO CAS9 | rCyp2E1 (1) | hCyp2E1 | F | 206 | n/a | 110 | 6.8% | 0.49 | n.s. | n.s. |
| 14 |  | NO CAS9 | rCYP3A1, r3A2 (2) | cis-hCYP3A4, Ins8 | F | 248 | 0, 0 | 132 | 6.5% | 0.47 | n.s. | n.s. |

**Table 3B:** Mouse AG gene conversion efficiency acting on cis AGC.

|  | cis-Mouse (8/25) |  |  |  |  | AG Gene Conversion |  |  |  |  |  |  |
| --- | --- | --- | --- | --- | --- | --- | --- | --- | --- | --- | --- | --- |
|  | A | B | C | D | E | F | G | H | I | J | K | K |
|  | Promoter-Cas9 | Cas9 Hetero or Homozygous | AG cassette gRNA target site (number of sites) | Donor chromosome Genetic Marker | Sex | Number Pups Examined | Intermarker Molecular Biology Events observed (B,C) | Number of Observed AGC Present, Including Molecu la (A+B+C) | % Gene conversion (AG Efficiency), Including Molecular | x2 p-val | % Gene Conversion Significance (n.s., p>0.05) | Interperated Amount of AG Observed ? |
| 1 | pSycp1-Cas9 R26 | Hetero | mFcgr2, mFcgr3 (2) | cis-hFcgr2, hFcgr3 | M | 200 | 0, 1 | 107 | 7.0% | 0.48 | n.s. | 0.3% |
| 2 |  | Homo | mFcgr2, mFcgr3 (2) | cis-hFcgr2, hFcgr3 | M | 62 | 0, 2 | 33 | 6.5% | 0.72 | n.s. | 1.6% |
| 3 |  | Hetero | mFcgr2, mFcgr3 (2) | cis-hFcgr2, hFcgr3 | F | 206 | 0, 0 | 108 | 4.9% | 0.62 | n.s. | n.s. |
| 4 | Sycp1-P2A-Cas9 | Hetero | mFcgr2, mFcgr3 (2) | cis-hFcgr2, hFcgr3 | M | 205 | 0, 7 | 103 | 0.5% | 0.96 | n.s. | 1.7% |
| 5 |  | Homo | mFcgr2, mFcgr3 (2) | cis-hFcgr2, hFcgr3 | M | 62 | 0, 4 | 32 | 3.2% | 0.86 | n.s. | 3.2% |
| 6 |  | Hetero | mFcgr2, mFcgr3 (2) | cis-hFcgr2, hFcgr3 | F | 242 | 1, 3 | 124 | 2.5% | 0.79 | n.s. | 0.8% |
| 7 |  | NO Cas9 | mFcgr2, mFcgr3 (2) | cis-hFcgr2, hFcgr3 | F | 211 | 0, 0 | 104 | -1.4% | 0.88 | n.s. | n.s. |
| 8 |  | NO Cas9 | mFcgr2, mFcgr3 (2) | cis-hFcgr2, hFcgr3 | M | 232 | 0, 0 | 125 | 7.8% | 0.40 | n.s. | n.s. |

**Table 3C:** Mouse AG gene conversion efficiency acting on trans AGC.

| trans-Mouse (8/25) |  |  |  |  | AG Gene Conversion |  |  |  |  |  |  |  |  |  |  |
| --- | --- | --- | --- | --- | --- | --- | --- | --- | --- | --- | --- | --- | --- | --- | --- |
|  | A | B | C | D | E | F | G | H | I | J | K | L | M | O | P |
|  | Promoter-Cas9 | Cas9 Hetero or Homozygous | AG cassette gRNA target site (number of sites) | Donor chromosome Genetic Marker | Sex | Number Pups Examined | Molecular Biology Events observed (C + D) | Number of Observed AGC Present, Including Molecular (A C + D) | Number of Observed AGC Present, Including Molecular (B + C + D) | % Gene conversion (AG Efficiency), Including Molecular (A + C + D) | % Gene conversion (AG Efficiency), Including Molecular (B + C+ D) | X2 p-value (A + C + D) | X2 p-value (B + C + D) | % Gene Conversion Significance (n.s., p>0.05) | Interperated Amount of AG Observed? |
| 1 | pSycp1-Cas9 R26 | Hetero | mFcgr1 (1) | trans-hFcgr1, Del24 | M | 219 | 4 | 111 | 112 | 1.4% | 2.3% | 0.886 | 0.811 | n.s. | 0.9% |
| 2 |  | Hetero | mFcgr1 (1) | trans-hFcgr1, Del24 | F | 48 | 0 | 22 | 26 | -8.3% | 8.3% | 0.683 | 0.683 | n.s. | 0 |
| 3 |  | Hetero | mFcgr2, mFcgr3 (2) | trans-hFcgr2, hFcgr3 | M | 182 | 6 | 91 | 97 | 0.0% | 6.6% | 1.000 | 0.53 | n.s. | 1.6% |
| 4 |  | Hetero | mFcgr2, mFcgr3 (2) | trans-hFcgr2, hFcgr3 | F | 130 | 2 | 57 | 75 | -12.3% | 15.4% | 0.321 | 0.21 | n.s. | 0.8% |
| 5 |  | Homo | mFcgr2, mFcgr3 (2) | trans-hFcgr2, hFcgr3 | F | 46 | 0 | 24 | 22 | 4.3% | -4.3% | 0.835 | 0.83 | n.s. | 0 |
| 6 | Sycp1-P2A-Cas9 | Hetero | mFcgr1 (1) | trans-hFcgr1, Del13 | M | 114 | 0 | 56 | 58 | -1.8% | 1.8% | 0.895 | 0.895 | n.s. | 0 |
| 7 |  | Hetero | mFcgr1 (1) | trans-hFcgr1, Del13 | F | 105 | 0 | 58 | 47 | 10.5% | -10.5% | 0.448 | 0.448 | n.s. | 0 |
| 8 |  | Hetero | mFcgr2, mFcgr3 (2) | trans-hFcgr2, hFcgr3 | M | 123 | 5 | 58 | 70 | -5.7% | 13.8% | 0.655 | 0.278 | n.s. | 2.0% |
| 9 |  | Homo | mFcgr2, mFcgr3 (2) | trans-hFcgr2, hFcgr3 | M | 203 | 6 | 99 | 110 | -2.5% | 8.4% | 0.804 | 0.399 | n.s. | 1.5% |
| 10 |  | Hetero | mFcgr2, mFcgr3 (2) | trans-hFcgr2, hFcgr3 | F | 78 | 0 | 37 | 41 | -5.1% | 5.1% | 0.749 | 0.749 | n.s. | 0 |
| 11 |  | NO Cas9 | mFcgr1 (1) | trans-hFcgr1, Del24 | F | 203 | 0 | 109 | 94 | 7.4% | -7.4% | 0.457 | 0.457 | n.s. | 0 |
| 12 |  | NO Cas9 | mFcgr2, mFcgr3 (2) | trans-hFcgr2, hFcgr3 | F | 200 | 0 | 91 | 109 | -9.0% | 9.0% | 0.368 | 0.368 | n.s. | 0 |
| 13 |  | NO Cas9 | mFcgr1 (1) | trans-hFcgr1, Del24 | M | 210 | 0 | 108 | 102 | 2.9% | -2.9% | 0.770 | 0.770 | n.s. | 0 |
| 14 |  | NO Cas9 | mFcgr2, mFcgr3 (2) | trans-hFcgr2, hFcgr3 | M | 187 | 0 | 94 | 93 | 0.5% | -0.5% | 0.959 | 0.959 | n.s. | 0 |

### Promoter

In the rat, the *Ddx4-*Cas9 promoter configuration provided higher gene conversion rates than the *Sycp1* promoter. Significant AG efficiency, 67%, occurred in the female homozygous *Ddx4-*Cas9 arrangement when paired with the r*Cyp3A1*-r*Cyp3A2* AGC. Homozygous Sycp1-Cas9 females paired with the r*Cyp3A1*-r*Cyp3A2* AGC afforded 22% AG efficiency, while a female strain with *rCyp2E1* target paired with the homozygous *Ddx4* driven Cas9 exhibited 20% AG efficiency in the homozygous configuration, which did not reach statistical significance based on sampling 125 F2 offspring (molecular confirmation was not possible for the *rCyp2E1* locus due to the absence of a nearby marker). In the rat promoter-AGC pairings just described, molecular detection supports the conclusion that gene conversion events occurred and are consistent with the robust super-Mendelian excess observed.

In the mouse, both Cas9 promoter configurations, with pSycp1 or Sycp1 drivers, provided low levels of AG gene conversion in the 0.8% - 3.2% range, as demonstrated using the molecular test. These strains targeted catalysis of gene conversion at the linked mFcgR2-mFcgR3 pair in cis or trans configurations, and the mFcgR1-trans Del 13/24 marked chromosomes (Table 3B, 3C).

The striking difference in efficiency in the rat versus mouse and the striking difference in the rates of detection with the Mendelian method versus the molecular method illuminate features of the technology related to the timing of Cas9 expression and level of Cas9 expression, and target site homology are discussed below.

### Parental sex

Female rats configured to express Cas9 from the *Ddx4* or the *Sycp1* promoter and an AGC encoded gRNA directed to the rat Cyp3A1 plus Cyp3A2 locus exhibited AG efficiencies ranging from 0.7% to 67% (Table 3A). Homozygous female *Sycp1* driven Cas9 operating with the *Cyp3A1-Cyp3A2* AGC provided 22% AG efficiency. Heterozygous female *Sycp1* strains provided low efficiency, but the strain exhibited molecularly detectable AG events at the 0.7% rate. In males, the *Ddx4* or *Sycp1* driven Cas9 arrangement working with the *Cyp3A1-Cyp3A2* locus AGC afforded 0.9% - 1.0% AG efficiency. Thus overall, female rat configurations were more successful than the corresponding male configurations.

In mice, AG efficiency with either the pSycp1 or Sycp1 promoter targeting cis or trans-Fcgr2-FcgR3 or FcgR1-Del13/24 provided lower levels of AG efficiency, in the 0.8% to 3.2% range (Table 3B, 3C). No trend in male versus female performance was apparent, however. Thus, the better performance trend observed in female rats compared to males did not hold for mice.

### Number of Cas9-gRNA target sites

Single gRNA configurations provided no significant or very low levels of gene conversion. For example, in the rat where only a single Cas9-gRNA target site was available, as in the 2E1-AGC situation, we detected no significant AG gene conversion. In the rat, the presence of two gRNA target sites sustained AG efficiencies ranging from 22% - 67% (Table 3A). Some multiple-cut site arrangements, however, drove only low levels of gene conversion with efficiencies in the 0.7% - 1% range. These events were only observable using the molecular biological assay. Examples of this outcome include the heterozygous male *Ddx4*-Cas9 or the heterozygous male or female *Sycp1*-Cas9 configurations (Table 3A).

In the mouse, only one configuration of single gRNA targeted loci supported AG; in the heterozygous pSycp1-trans-FcgR1-Del24 configuration, we observed 0.9% AG efficiency. Three other single-site formats provided no detectable AG efficiency. In the mouse, the two gRNA cut site configurations were more successful; 9 of 12 such targets achieved AG efficiency in the 0.8-3.2% range (Table 3B, 3C). Thus, as in the rat, two gRNA cut sites in the mouse drove better than single site arrangements.

### Zygosity

More Cas9 activity is often associated with higher gene conversion frequencies. For example, high levels of AG and high levels of Cas9-gRNA associated indels were observed in the female rat *Ddx4* and *Sycp1* driven Cas9 expression strains operating on the *3A1-3A2* AGC (Table 3A and Table SR9A). All four homozygous vs. heterozygous strain pairs in the rat displayed higher levels of gene conversion in the homozygous versus heterozygous strains. The difference was often striking, with little gene conversion in the heterozygous state but significant conversion in the homozygous state. Greater frequencies of indels were also observed in the homozygous versus the heterozygous state. There was no obvious trend for homozygous strains in the mouse performing gene conversion more efficiently than heterozygous strains. In the mouse configurations tested, the AG rates are low for both hetero-and homozygous mice, thus the effect may be obscured by the small signal. Therefore, additional Cas9 provided by the homozygous configuration in rats, but not mice, results in more frequent chromosomal hydrolysis and gene conversion.

### The importance of the level of Cas9 activity for AG efficiency

Our comprehensive measurement of the number of rat offspring with a target site indels allowed us to investigate the relationship between Cas9-induced DSBs, indels, and AG gene conversion efficiency. We interpret the indel formation rate as a reflection of the gRNA’s DNA targeting ability and the level of Cas9 expression in the gamete and embryo. Figure SR12 shows the correlation for 14 configurations divided into two classes, AGC encoded gRNAs with two hydrolysis targets and those gRNAs with a single target. The correlation between indels and AG efficiency is modest for AGC encoding gRNAs with one cut site R^2^=0.49. The correlation for AGCs encoding gRNAs with two chromosomal targets is much stronger, R^2^=0.65. The strength of this correlation, considering the hundreds of steps and proteins involved in gene conversion and meiosis, emphasizes the important contribution of Cas9 expression level and its associated DSB to promote efficient gene conversion.

In the mouse, the lower levels of AG efficiency observed do not permit us to assess similar correlations between indel occurrence and AG occurrence.

### Absence of Cas9

There is no background rate of EJ repair and AG gene conversion in rats and mice. Our studies of Cas9-induced indels studied 454 F2 offspring rats with two different AGC encoded gRNA configurations, but in which Cas9 was absent. We detect no EJ repaired indels at the gRNA target site in these strains, including exceptionally successful cis 3A1/3A2 AG configuration (Table SR9A). Similar tests in mice examining 1243 F2 offspring from animals with six different AGC encoded gRNA configurations did not detect a single inde (Table SR9B). Thus, the presence of EJ indels requires Cas9 and gRNA to be present; there is no background rate of EJ repair. This observation points to the absolute requirement for Cas9 caused DSB to cause both the indels and, as discussed below, the similar finding for AG gene conversion in rats and mice.

### Rare large deletions

We observed some rare genotypes inconsistent with expected Mendelian or gene conversion outcomes in the rat 3A1/3A2 bearing strains. We investigated these events using a PCR-based assay to detect the deletion of regions between the two gRNA target sites Figure SM1. Such large deletions (LD) might occur when two gRNAs-Cas9 induced DSBs are sealed by an EJ reaction or another repair mechanism. In addition to the confirmed large deletions, our assay detected a few genotypes consistent with very large deletions, extending beyond 55 kb. In the rat AG configurations encoding two gRNAs, observed 95.7% of the genetics outcome are consistent with Mendelian expectations or standard gene conversion outcomes (Table SR10). Large deletion events occurred in 79/1871 offspring, a rate of 4.2% of the total 3A1/3A2 offspring. The majority were repaired by exact re-ligation at the gRNA cleavage sites. These deletion events occur at a rate of 25.1% of the rate at which AG occurs in the 3A1/3A2 rats. These deletions reveal that two Cas9 targets, a substantial distance apart, can be re-ligated precisely by end joining. Given our desired outcome, Cas9-induced gene conversion, these events represent an unproductive competing pathway. In the mouse, we saw no genotypic outcomes that were inconsistent with either Mendelian or active genetics outcomes (Table SR10). However, the genotype D of figure 1 could have a LD that would not be detected by our standard genotype assay. Therefore, in the mouse, we developed LD assays for all the cis and trans two gRNA mouse configurations. We conducted these tests in five trans-configured offspring with genotype D, and 116 of cis-configured offspring with genotype D. No mouse events were detected. Thus. large deletions are a feature of the Rat 3A1/3A2 configuration and not of any of the mouse configurations. We believe that this observation is likely due to the extremely high homology in the 3A1/3A2 test case.

## Discussion

Our efforts to develop broadly applicable Active Genetic systems in mammals that are highly efficient, robust, and reusable have made progress. We have improved gene conversion efficiency in the rodent from the previous *reusable* best of 5.5% to our best of 67% in females, a 12-fold improvement over the best *reusable* mouse finding. The results provide new information about the configurations and requirements for establishing robust and efficient gene conversion systems in rodents.

### Cellular localization and biochemical activity of Cas9

The cellular location and timing of Cas9 expressed from different promoters have emerged as key elements in developing highly efficient gene conversion systems (Grunwald et al., 2019; Weitzel et al., 2021; Terradas et al., 2021; Bier, 2022). We prepared two rat AG accelerator strains: the Ddx4-driven Cas9 strain (Vasa in Drosophila and humans, and often called Mvh in mice) and the Sycp1-driven Cas9 strain. Likewise, mice with Sycp1 or pSycp1-Cas9-driven Cas9 configurations were also built (pSycp1-Cas9 a strain using a promoter fragment of the Sycp1 promoter).

Both rat and mouse strains express readily detectable levels of Cas9 protein in the developing sperm lineage, including cells with the size, shape, and position of zygotene-pachytene stage and cells with 4N DNA content. In these sperm progenitor cells, nuclear localization of Cas9 is visible, consistent with the presence of a nuclear localization sequence on the encoded Cas9 transgene. These findings indicate that we achieved our objective of targeting Cas9 expression to the nucleus of primary spermatocytes with 4N DNA content and leptotene-through-pachytene stage sperm cells as planned.

The Cas9 expressed in the different rat configurations is biochemically active, as revealed by the presence of EJ repaired indels at the gRNA target sites in isolated testis cells from strains bearing Cas9 and AGC transgenes. In addition, 12 of 12 rat strains and 16 of 16 mouse strains produced EJ repaired indels when configured to express Cas9 in the presence of an AGC. These tests include both male and female Cas9-bearing parents. Most rat and mouse strains generated indels and ACG conversion events consistent with Cas9 activity in the gamete or at the one cell-post fertilization stage of embryogenesis and other stages. Thus, we detected indels in the germlines of female and male rats or mice parents carrying Cas9-AGC transgenes, consistent with Cas9 biochemical activity during gametogenesis.

### What configurations allow active genetics in the rat or mouse?

We varied five key parameters to provide a better understanding of arrangements promoting AG gene conversion. Thus, we investigated the promoter for Cas9, the number of potential target sites, the zygosity of Cas9, the sex in which AG might occur, and the cis or trans configuration.

In rats, examining 12 configurations, we found three of the twelve tested configurations supported efficient AG occurring at rates ranging from 22-67%. These events were detected both by super-Mendelian excess and as 42 molecularly detected AG events out of 895 pups examined. In the four other rat configurations, the molecular detection method identified 17 AG gene conversions out of 976 pups examined, implying 0.7%-1% efficiencies. Super-Mendelian excess did not reach statistical significance in these four cases by examining 216 or more F2 offspring.

In mice, we investigated 16 Cas9 bearing configurations varying the five key parameters to provide a better understanding of the requirements for efficient AG gene conversion. In none of the 16 configurations was statistically significant gene conversion observed by Super-Mendelian excess. In most of the 16 configurations, however, we did observe molecularly detected AG gene conversion events. We observed 41 molecular marker detectable events out of 2530 pups examined, with efficiencies ranging from 0.8%-3.2%.

### Absence of Cas9

In the absence of Cas9 expression in the rat and in separate lines containing two different AGCs, zero out of 454 (0/454) pups examined showed any evidence of AG by the molecular detection assays or by Mendelian excess. In the presence of Cas9 and the same two AGCs, 59 out of 2811 rat offspring exhibited AG events in the molecular detection assay.

In mice, we examined 3507 F2 offspring using three AGC; in the *presence* of Cas9, 41 of 3507 offspring exhibited AG events by the molecular detection assay. In the *absence* of Cas9, we saw zero AG events in 1243 F2 Cas9 mice (0/1243) using the molecular detection assay, and these showed no excess above the Mendelian expectation. We conclude that there is no detectable background rate of AG in the absence of Cas9 in the mouse.

The complete lack of molecularly detected events in the absence of Cas9 in both the rat and mouse and the consistent detection of molecularly detected events in the presence of Cas9 points to the significant acceleration of gene conversion near a Cas9-induced DSB site. These findings are consistent with the requirement for a DSB for HDR observed in species including yeast, *Drosophila*, plants, and rodents has been extensively studied (Zickler and Kleckner, 2015; Hunter, 2015; Kockler et al., 2021; Xue and Greene, 2021).

### Comparison to other findings

The observed AG frequencies observed in this study are superior to reusable mouse configurations using the Spo11 promoter, synthetic promoters, or the mouse Vasa promoter (Grunwald et al., 2019; Pfitzner et al., 2020; Weitzel et al., 2021). Those mouse studies reported AG efficiencies ranging from 0% to 5.5% when applied in the Tyrosinase locus in a trans-configured single gRNA setting. In a single-use and indirect, Cre-Lox configuration, the Vasa promoter afforded 44% AG efficiency in female mice and 0% efficiency in male mice. Another similarly configured single-use Cre-Lox-Stra8 promoter produces no AG events in either male or female mice (Grunwald et al., 2019). Our reusable Ddx4 rat construct provides exceptionally high AG efficiency in some cases.

In contrast, Pfitzner et al. made a different construct in which a Ddx4 promoter fragment was placed ahead of Cas9 and integrated the construct randomly in the genome. This promoter fragment-driven Cas9 failed to provide any gene conversion. The Ddx4 construct used here placed Cas9 behind a peptide cleavage site fused to the C-terminus of the Ddx4(Vasa) protein. In favorable cases, it provided gene conversion efficiency in the 55-67% range. The configuration used here allowed the natural expression of Ddx4 and induced Cas9 expression at the same cellular stage as DDX4 during meiosis Prophase I in primary spermatocytes and likely in a similar stage in the developing egg. The difference in the performance of the fused Ddx4 arrangement versus the promoter fragment arrangement points to the importance of producing sufficient levels of Cas9 expression at developmental times similar to the naturally expressed DdX4 protein. A Sycp1-Cas9 configuration also provided 22% gene conversion efficiency, so while Ddx4-Cas9 exhibited the highest AG efficiency, other promoter configurations may also sustain adequate performance.

The Ddx4 or Sycp1-driven Cas9 configuration may not be the only element contributing to the exceptional efficiency observed in rats. The cis configuration of the r3A1/3A2 target locus is also important. This target element has unique features not found in literature reports or our other rat or mouse configurations. First, this configuration has two gRNA cut sites; all other target sites reported in other studies have only a single gRNA target. Second, the 3A1 and 3A2 target sites are nearly identical for up to 200 bases on either side of the gRNA cleavage sites. In our test of other mouse strains with two gRNA targets, no such homology was present, and the observed gene conversion rates were substantially lower, in the 0.8-3.2% range. Together, these observations highlight the importance of deploying the right promoter to drive sufficient Cas9 expression at the right stage of meiosis. They emphasize the importance of employing two gRNA cut sites. In addition, our findings uncover a benefit of high homology around the two gRNA cut sites, which is a novel observation in the field.

### Seven informative design features and outcomes

We can extract seven instructive design features from our study which may illuminate key mechanisms underlying efficient gene conversion events. 1) Promotion of gene conversion is enhanced using two gRNA target sites. 2) Cas9 Expression during meiosis prophase I improve AG efficiency. 3) Homology near the Cas9 gRNA improves AG efficiency. 4) Long-range gene conversion tracks occur and have precedent. 5) Males display less AG than females. 6) Increased Cas9 levels stimulate greater AG. 7) High 22-67% AG gene conversion efficiency was detected by Mendelian excess, but, in the same configurations, only a 2.3% conversion rate were observed by the molecular detection method.

1. *Two gRNA target sites versus one:* Our data reveal a consistent trend toward improved AG gene conversion efficiency in settings with two gRNA-Cas9 target sites rather than only one. In the rat, two gRNA configurations were the only configuration that provided highly efficient gene conversion; the single gRNA configurations were unsuccessful. We note that not all two gRNA configurations exhibited high-efficiency AG gene conversion, so other factors also contribute to efficiency. In the mouse, 3 of the 4 cases wherein no molecular conversion events were detected were single gRNA arrangements. In contrast, in the mouse, 10 out of 12 two gRNA configurations resulted in molecularly detectable AG events. Thus, there appears to be a trend wherein two Cas9-gRNA target sites improve efficiency. We speculate that two cut sites may create a large gap that is repaired using the homologous chromosome or the sister chromosome in the leptotene-to-pachytene gametocyte. We see parallels to a similar form of cooperativity between dual gRNAs versus single gRNA configurations observed for so-called Element Reversing the Autocatalytic Chain Reaction (ERACR) elements that delete and replace gene-drive elements in Drosophila (Xu et al., 2020). In addition, we speculate that cutting the genome at two potential sites improves the frequency of Cas9-mediated DSB formation and thus increases the frequency of initiation of HDR.
2. *Cas9 Expression during Meiosis prophase I improves AG efficiency:* Our most successful AG configuration is the DDx4-Cas9 arrangement. This strain expresses Cas9 in primary spermatocytes, cells undergoing meiosis I. These cells are in the leptotene to pachytene stage when homology-directed repair is highly active. Our second most successful strain is the homozygous Sycp1-driven Cas9 strain. This strain expresses high levels of Cas9 in spermatogonia. In testis, 7.5% of the rat and 2.2% of the mouse 4N DNA content cells, cells in leptotene-pachytene, stain positive for the GFP surrogate marker indicating that there is significant Cas9 expression in the target cell population. Given highly efficient AG in female parents, we presume that the sperm findings and our observed female AG rates indicate similar meiosis prophase I targeting in the developing primordial oocyte.
3. *Homology near the Cas9 gRNA target site:* Why is the *Cyp3A1/3A2* AGC configuration better than the other ACG configurations? The 3A1/3A2 arrangement has a unique feature; the 3A1 and 3A2 targets sites are nearly identical for up to 200 bases around the site, including perfect identity for the first 20 bases on either side of the target site, and only 5 miss-matches in the 50 bases on either side of the target site. The other two-cut case, the Mouse *FcgR2b/FcgR3* AGC, has no such homology near the target sites, with 27 miss-matches in the 40 bases on either side of the cleavage site. This high degree of local sequence identity may be significant. The long-track gene conversion process is highly favored when the homology at the DSB site between the donor chromosome and the recipient is very high, for about 200 bp (Stewart et al., 2021). We speculate that having an excellent match at both ends of the DSB allows for two sites of initiation of D-loop invasion and increases the probability of starting and completing the repair leading to more successful BID and our observed high efficiency in the *Cyp3A1/3A2* configuration The high homology may also provide a stable D-loop, leading to higher repair completion rates (Hunter and Kleckner, 2001; Roy and Greene, 2020). However, the precise mechanism by which the near-perfect homology at two distantly located sites may promote highly efficient gene conversion needs further investigation to test this hypothesis.
4. *Length of Gene conversion:* The rat cis-*3A1/3A2* configuration provided highly efficient gene conversion of a chromosomal region of more than 55kb, observed by super-Mendelian excess. The mouse trans arrangement of FcgR2/FcgR3, spaced 83 kb apart, provided lower frequencies of molecularly detectable AG. However, the mouse events were all intermarker events, with no overall deviation from Mendelian transmission, implying few events occurred outside of the markers. Long track gene conversion, a break-induced repair terminated by a crossover or a crossover-like event, is a likely mechanism behind gene conversion covering such long stretches (Kockler et al., 2021). Long track gene conversion has been observed in diploid *Saccharomyces cerevisiae* encompassing chromosomal regions from 6-464 kb (Stewart et al. 2021). In mammalian cells, long-tract gene conversion has been observed during mitosis, encompassing a minimum of 11 kb (Neuwirth et at. 2007). In other mammalian settings, long-tract gene conversion has also been observed (Scully et al., 2019). Thus, the observation of gene conversion in regions as large as 55kb has precedent.
5. *Male vs. female:* HDR terminated by crossover is our postulated mechanism by which gene conversion occurs. Thus, components of the crossover machinery are likely to contribute to this process. For example, a crossover chiasma in one location could block another event for many megabases in the surrounding region. Crossover interference is more pronounced in male mice and rats by about 30% (Petkov et al., 2007; Jensen-Seaman et al., 2004). This known phenomenon led us to speculate that females will usually have higher efficiency than males. In the rat, the AG levels observed in males were lower than the best performance in females for each pair of otherwise identical arrangements tested, which is similar to mouse findings at the Tyrosinase locus (Weitzel-Cooper, 2020). Other features of AG in the male may also contribute, including the lack of synchrony, the chromosomal compaction during spermatogenesis, and the concentration of Cas9 and gRNA in the developing sperm.
6. *Increasing Cas9 levels results in greater AG:* More Cas9 activity was often associated with higher frequencies of AG. All four of the Cas9 homozygous-heterozygous strain pairs in the rat provided more gene conversion than the heterozygous strains. Often the difference was striking, with just 0.7% gene conversion in the heterozygous state but a 22% conversion in the homozygous state, as in the female Scyp1-Cas9 rat. This dramatic dose-dependent difference in efficiency suggests that a threshold level of Cas9 expression may need to be exceeded for the process to be highly efficient. Our data show a correlation between the number of indels and AG gene conversion efficiency in the rat. We plotted the level of gene conversion activity vs. the level of Cas9-induced activity as assessed by the percentage of F2 animals with indels in the twelve strains and two control strains tested (Figure SR12). When two gRNA target sites are available, the linear model R^2^is a moderately strong 0.65 (N=8). When one gRNA target site is available, the R^2^ is weaker 0.49 (N=6). These findings underline the value of having more than one gRNA site near the target gene conversion location(s). Given the complexity of the gene conversion pathways (Kockler et al., 2021), meiosis (Zickler and Kleckner, 2015; Bolcun-Filas and Handel, 2018), and gamete development (Ernest et al., 2019; Wang and Pepling; 2021; Zamboni et al., 1972), processes involving thousands of proteins and hundreds of steps, the strength of these correlations is remarkable. They reinforce the conclusion that timing and levels of Cas9 activity in the developing gamete are critical. The correlation between Cas9 level and AG efficiency in the mouse is not strong as in the rat, we believe that this low correlation is driven by the low levels of mouse AG efficiency seen, in the 0-3% range. In this situation observation of a correlation is obscured by the small signal size. For the same reason, in the mouse, Cas9 homozygous configurations did not increase copying efficiency in paired heterozygous versus homozygous strains. The trans mFcgR2/mFcgR3 AGC in the presence of either the pSYCP1 or SYCP1 driven Cas9 accelerator construct are examples of these findings. The contrast between the rat and mouse may be due to the special high homology property of the 3A1/3A2 configuration, and its resulting high AG efficiency, rather than reflecting a species difference. In the mouse, mFcgR2/mFcgR3 pair lack homology at the gRNA cut sites. We believe that the homology in the 3A1/3A2 setting is the dominant factor and that level of Cas9 expression may be a second contributing factor.
7. *Super-Mendelian Excess vs. Molecular Detection:* We observed very low levels of molecularly detectable independent AG events, 1-3%, but much higher rates of super-Mendelian excess, up to 67% in several cases. For example, in the heterozygous female rat Ddx4-Cas9 strain operating on the cis 3A1/3A2 AGC, we can see an imputed 145 Mendelian excess events out of 526 pups examined, whereas, with the molecular assay in the same setting, we see just 21 events in the 526 pups. These observations could be explained by gene conversion having a strong overlap with the mechanism of meiotic crossover and crossover interference. The process may employ the Homology Directed Repair mechanism terminated by a crossover. The molecular test can only detect an AG event if a crossover happens between the two markers. Our molecular markers vary between 1.7 kb and 83 kb (Figure 2) from the Cas9 target site; thus, we expect crossover events between these markers to be rare by the usual centimorgan standard – on the order of 0.08%. This line of reasoning, coupled with the much higher gene conversion rate seen by the Mendelian excess test, leads to the understanding that most gene conversion events must occur outside our genetic markers. They could occur long distances from the gRNA-Cas9 target, maybe hundreds-of-thousands bases distant. The DSB formed by Cas9-gRNA accelerates gene conversion but does not supersede the need for a crossover to terminate the synthetic step. Therefore, the rules of crossover and crossover interference constrain short-distance outcomes, consistent with our observations.

### Mechanisms for gene conversion during AG

We postulate that the homology directed repair process initiated by the Cas9-induced DSB is likely be terminated by crossover events. In our data we find parallels to previous observations of long tract gene conversion events (Stewart et al., 2021; Neuwirth et at. 2007; Scully et al., 2019). Long tract conversion is a form of Synthesis dependent strand annealing (SDSA) or break-induced repair (BIR) and may be terminated by crossover or the end of the chromosome (Kockler et al., 2021). A crossover is a trade of one chromosome piece for another; hence it does not add alleles and is neutral with regard to gene frequency. There must be a chromosome homolog-biased repair step to observe an increase in the number of AG markers. Therefore, crossover alone cannot explain the loss of heterozygosity in strains with 20-60% AG efficiency. Long tract conversion events based on Synthesis dependent strand annealing (SDSA) or break-induced homology-directed repair (BIR) are candidate pathways for these outcomes (Kockler et al., 2021; Stewart et al., 2012). Our findings are consistent with large-scale events, as large as 55 kb, such long-distance repair events have been documented in yeast, and reported in mammalian cells (Stewart et al., 2012; Neuwrith et al., 2007).

## Conclusions

In this study, we assess the gene-conversion efficiencies of various AG configurations in both rats and mice. These findings reveal that AG-mediated gene conversion is widely applicable to many genes and points to the complex interaction of multiple variables that are key to optimizing the performance of these mammalian AG systems. Four variables are important. The promoter is critical, and the timing of expression to the leptotene to pachytene stage of meiosis prophase-I is important. Temporal and quantitative promoter features are also important as they must drive sufficient levels of Cas9 expression at the appropriate time during early meiosis. Special features of the particular Cas9-gRNA target sites, including extended homology between pairs of gRNA cut sites, appear to be as critical as the promoter to drive very high levels of gene conversion. Two gRNA cut sites substantially improve AG gene conversion efficiency. Finally, the sex of the parents in which the gametes form also affects AG, with females being generally more efficient than males.

We have identified several previously untested promoters in mice and rats that foster high AG efficiency. We place these in order of their ability to make biochemically active Cas9 and to promote AG gene conversion as follows: Ddx4 in rats and the Sycp1 promoter in rats and mice, then the pSycp1 promoter in mice. Our findings suggest a strong correlation between AG efficiency and the level of Cas9 expression. Our data uncovered the unexpected benefits of high homology at two gRNA cut sites as a key element of efficient AG. These studies provide new data which may guide future efforts to develop practically useful strategies.

We suggest several steps to continue these investigations. First, we recommend that two or more gRNAs be employed in future tests. We speculate that intra-Cas9 target distance should be explored, and those explorations should include distances as large as the crossover interference distances, 62.5-95 Mb in mice. Given the limited target distances available on small chromosomes, we would place the ACG on chromosomes 1-5, the biggest chromosomes. Finally, other promoters should be investigated.

To conduct further investigations and to develop an accelerator strain(s) with the capability of robust application, a special set of sites may need to be placed onto one, or several, of the larger chromosomes. For example, this special accelerator strain could have an array of engineered target loci sites incorporating high homology at two target loci and an optimized distance between the loci. Additionally, further steps should be undertaken to optimize the timing and strength of promoters driving Cas9 expression. With combinations of these changes, we envision developing generalizable AG systems with robust and reliable performance characteristics required to generate complex model genetic systems in mammals.

## Acknowledgments

Synbal would like to thank Drs. Ethan Bier and Kim Cooper for their helpful discussions. Drs. Stephen Hedrick and Rick Morrissey provided valuable input. The interest and support of Drs. Ho Cho, Kate Blease, Rama Narla, and Winston Thomas are gratefully acknowledged. Synbal would like to thank Grant Bolgard and Márton Münz of The Bioinformatics CRO for their assistance with bioinformatics analysis. We also thank Dr. Chris Premanandan, DVM, PhD, ACVP, ACT, Ohio State Univ., for providing histopathology assessments. TransViragen would like to acknowledge the excellent technical support of the staff of TransViragen and the University of North Carolina Animal Models Core Facility.

## Supplemental Methods

### Indel calculations

Tables SR9 report data indels that took place at two loci and are reported as the number of events at locus one followed by locus two. These figures report summary statistics which are calculated as the average number of events at either locus, rounded up to the next whole integer. Indel% =Number of observed indels at locus/total number of F2 pups tested. We have scored the point in cell development in which an Indel occurs based on the combination of the Taqman® results and the ICE algorithm estimated indel frequencies (Hsiau et al. 2018) using the following metrics: One cell Post Fertilization stage cut (column G of Figure SR9), AGC Present, and 100% of a single indel, or AGC Absent and 100% of a single indel; Gamete Stage Cut or One Cell stage cut(column H of Figure SR9), AGC Absent, with 50% single indel and 50% WT; and, Two or more Cell Stage Cuts (column I of Figure SR9), AGC Present or Absent with WT at ∼75% or ∼50%, and <35% of one or more indels, indels summing to about 25% or 50%.

### Active Genetic gene conversion efficiency calculations

In the cis configuration, we measured the AGC count for both loci independently as locus one, followed by locus two. These numbers may not be identical due to differences in the efficiency at which the gRNA cuts each locus and chromosomal context factors in the intermarker region. AG efficiency includes AGC counts plus molecularly detected counts. In the cis configuration (Figure 3A or 3B), events at either marker locus are independent AG events; in the cis setting, we calculated total AG events as AGC count (Type A, Figure 1) plus the sum of the molecularly determined events (Type B or C, Figure 1). In the trans configuration, locus 1 or 2 (Figure 1) are on different chromosomes; thus, each could independently be the target of gene conversion. Therefore, we report (Figure 3C) the AG efficiency at locus 1 and 2 separately. Like the cis case, we use the sum of the molecularly determined events plus the AGC count at each locus. AG efficiency (Figure 3A, 3B, or 3C) is calculated as AG Eff% = (((number of AGC positive F2 pups observed/number of F2 pups tested)*100)-50)/50. Thus 100% efficiency indicates that all the gametes produced in the F1 parents contained the AGC, and 0% efficiency indicates that 50% (i.e., the Mendelian expectation) of the gametes produced in the F1 parents have the AGC. A Chi-squared test is conducted and judged significant if the AG Eff% imputes a p<0.05, the AG Eff% is placed in the interpretation column. If the value is not statistically significant, the AG% is calculated as the sum of molecular events/the number of pups examined/2, which is reported as the interpreted amount of AG. In cases where zero molecular events were observed or molecular events cannot be measured, and p>0.05, not significant is reported, n.s.

### Statistical considerations

How many meiosis events and offspring must we examine to detect Active Genetics occurring at various efficiencies? We used two methods for detecting Active Genetic events, molecular detection and the excess over the Mendelian expectation. Molecular detection provides an unequivocal demonstration of Active Genetic events but is limited to detecting events with an intermarker crossover. However, detection by Mendelian excess offers an opportunity to examine loci with no conveniently close DNA sequence markers. It allows the detection of large-scale events, including events outside genomic markers; thus, it is not biased by the location of convenient markers. Mendelian excess detection uses a Chi-squared test to assess the potential excesses of a locus versus the Mendelian expectation of 50%. These Chi-squared tests used a right-tailed test with one degree of freedom and a significance level of 0.05. A power calculation based on proportions using alpha =0.05 and 1-beta = 0.95 yields the conclusion that to detect Active Genetics at 10% efficiency; the test must examine 900 offspring at 25% efficiency, 195 offspring at 30% efficiency, 88 offspring, and at 60% efficiency, 24 offspring. (Proportion power tool implemented by HyLown Consulting LLC; 2019 https://powerandsamplesize.com/Calculators/Test-1-Proportion/1-Sample-Equality). We chose to examine efficiency to a lower bound of 25% and thus examined about 200 offspring from most F2 crosses.

Finally, another characteristic of robustness requires the process to happen in approximately equal frequency in all accelerator animals. Thus, in cases where high Mendelian excess was observed, we aimed to examine 5-10 parents and a 2-10 litters/parent to characterize the inter-animal variability of the process (Table SR11).

### Identification of appropriate promoters

We sought promoters that might provide an Active Genetic gene conversion system with improved performance in terms of efficiency and robustness. We investigated the literature searching for promoters restricted in expression to times around meiosis prophase I, with expression peaking in the leptotene-zygotene phase (LZ) and low in 2C spermatogonia. There are no genome-wide data on primordial oogonia; however, a thorough and wide-ranging study of spermatogonia provided guidance (DaCruz et al., 2016). In this study, transcriptomics in testicular cells separated by DNA content and then characterized by next-generation sequencing identified leptotene-zygotene and pachytene restricted gene expression. We sought genes with high LZ/2C expression ratios and lower expression levels in the 2C stage. We also focused on genes with prophase dominant expression with known involvement in chromosomal paring or synaptonemal complexes and recombination (Bolcun-Filas and Handel, 2018). The candidate genes *Sycp1, SYCP2, SYCP3, SYCE1, RAD51, DMC1, MSH4, MSH5*, and *HFM1* had LZ/2C expression ratios from 3 to 43. Based on literature findings, these proteins and promoters were further investigated regarding their known restriction to spermatogonia and oogonial cells, the degree to which the promoter was characterized, embryonic lethality, and sterility in males and females. We eliminated proteins that had expression outside of the testis and ovary, or which knockouts are embryonic lethal and for which the promoters were poorly characterized. These investigations led us to *Syp1* or *Sycp3* as promoters for further study. The LZ/2C ratio for *Sycp1* is 24, and *Sycp3* is 43 (DaCruz et al., 2016). The levels of expression of *Sycp3* mRNA are slightly higher in 2C than for *Sycp1*. *Sycp1*’s promoter has been studied in detail (Sage et al.,1999). *Sycp1* expression peaks at pachytene in oocytes and decreases rapidly afterward (Parades et al., 2005). Thus, we selected *Sycp1* as a developmentally regulated promoter for study as an Active Genetic candidate. *Sycp1* expression peaks during leptotene/zygotene and is the main transverse element of the synaptonemal complex (Bolcun-Filas and Handel, 2018). It is not expressed outside of gametogonigal cells, and its promoter has been studied (Sage et al., 1999). Sage et al. showed that a short 824 bp promoter could confer the expression of a reporter gene in primary spermatocytes in transgenic mice; however, the whole promoter is needed to cause expression in the embryonic ovary (Sage et al., 1999). This promoter fragment was selected since efficient male-active genetics is a goal. The Vasa promoter (*Ddx4* in the rat and human, and *Mvh* in the mouse) was also chosen for study because, in Drosophila, this promoter had functioned to promote efficient Active Genetics (Gantz, Bier, 2015). This gene has LZ/2C ratio of 13 and is restricted in expression to gametogonia cells during meiosis (DaCruz et al., 2016; Toyooka, 2000). During our studies, an unsuccessful mouse AG attempt using the Vasa promoter was published (Pfitzner et al., 2020); thus, a mouse *Vasa* construct was abandoned; however, our rat *Vasa* work was well advanced and continued^7^.

### Preparation of Cas9 Accelerator strains

Figure SM4 shows the configuration of six Cas9 accelerator strains produced during this work. Figure SM4 shows the configuration of humanizing minigenes and active genetic cassettes. Targeted transgenesis was achieved using the pronuclear injection method and assisted by gRNA-Cas9 targeting (Ittner and Gotz, 2007; Li, Li, et al., 2013; Wang et al., 2013). All humanizing cDNAs were constructed with the sequence found in the human genomic database. Lab-specific variations included the following steps and concentrations. Cas9 guide RNAs were designed using Benchling™ design tools. The selected guide RNAs were cloned into a T7 promoter vector, then T7 in vitro transcription (IVT) was conducted, followed by spin column purification, with elution in microinjection buffer (5 mM Tris-HCl pH 7.5, 0.1 mM EDTA).

In some cases, synthetic crRNAand tracrRNA (IDT, Coralville, Iowa) were redissolved in nuclease-free duplex buffer and annealed. Synthetic and T7 in vitro transcribed guide RNAs were pretested by transfection into rat or mouse embryonic fibroblasts (Yu et al., 2016) and assessed for functional efficiency using the T7-endonuclease 1 assay method, Alt-R Genome Editing Detection Kit (IDT, Coralville, IA) (Li, et al., 2013; Marshal et al., 1995; Sentmanat et al., 2018) or PCR followed by ICE analysis (Hsiau et al., 2018). In most cases, the gRNA providing the most cleavage was selected from 2-3 options. A supercoiled donor plasmid with ∼1kb homology arms flanking the region to be inserted was prepared. Then concentrated stocks were prepared by Qiagen High-Speed Maxiprep protocol (Qiagen, Germantown, MD) and resuspended in microinjection buffer. Embryo Microinjection: C57BL/6J zygotes were microinjected with 400 nM Cas9 protein, 25 ng/ul IVT guide RNA or 600 nM synthetic guide RNA, and 20 ng/ul donor plasmid in microinjection buffer. In some cases, concentrations were reduced by 50%. Taconic Sprague Dawley zygotes were injected with 50% of mouse reagent concentrations. Injected embryos were implanted in recipient pseudopregnant females. The pups were screened by PCR for the presence of the desired insertion events. TransViragen, Chapel Hill, NC, conducted the steps after plasmid cloning. Transgenic animals were detected using a series of five PCR reactions designed to detect the humanizing insert (A), the borders of the insert at 5’ and 3’ edge of the homology arms into the plasmid backbone (B & C), guarding against tandem integration events, and the 5’ and 3’ integration in the genome, through the homology arms, and into the insert (D & E). Animals with positive PCR results for PCR products A, D, and E and negative PCR products for the backbone (B & C) were likely candidate founders. These founders were bred to wild-type C57BL/6J or Sprague Dawley animals to transmit the targeting events and validate heterozygous animals. Each transgenic line was fully sequenced from ∼500 kb into the genomic DNA outside the homology arms through the entire insert sequence. Only those animals with sequences perfectly matching the planned sequence of donor plasmid DNA were carried forward for further breeding. During the breeding, non-target chromosome integrations were assessed by tracking that the offspring followed Mendelian expectations, demonstrating only one chromosomal copy of each transgene. We sequence the top five predicted off-target sites and checked those sites for the presence of indels. Prediction of the top five most likely off-target sites used the IDT’s CRISPR-Cas9 design checker tool (https://www.idtdna.com/site/order/designtool/index/CRISPR_SEQUENCE). All strains were free of indels at these sites. Thus, each animal has only a single copy of the target construct in the planned chromosomal location, and each insert and animal contain the planned genetic sequence.

### Active Genetic configuration

Figure SM4 shows the details of each rodent construct. Each rodent target locus was engineered to express the human homolog cDNA in exon-1 of the corresponding rodent gene. Each cDNA was arranged with a small SV40 artificial intron of 97 bp at the point in the cDNA where the human intron 1 started – such introns have been reported to improve transcription efficiency. The remainder of the minigene was the human cDNA. In many cases, we included a P2A or T2A self-cleavage sequence and the GFP sequence after the cDNA to facilitate protein expression analysis. The human 3’untranslated region and the poly-A+ site was included, followed by a strong three-frame stop to prevent unplanned read-through. Three-prime to the minigene, and in the opposite transcriptional sense, was an active genetic cassette that contains a multi-tissue U6 type III RNA polymerase III promoter followed by the gRNA needed to target the wild-type rodent sequence at the target site.

### Configuration of Rat Active Genetic Target Strain (Figure SM4)

The *rat Cyp2E1* was humanized by substituting the *hCYP2E1* minigene. The rat *Cyp2E1* locus has no convenient SNP nearby; therefore, we used genetic excess versus the Mendelian expectation to study Active Genetics at this locus. The rat homolog of human *CYP3A4* is rat *Cyp3A1,* and it is 83.2 kb from the very highly homologous gene, *rCyp3A2*. We humanized the *rCyp3A1* locus with a *hCYP3A4* minigene. Because of the very high homology between the *rCyp3A1* and *rCyp3A2* genes, our gRNA target cut twice during the humanization, humanizing the *3A1* locus with *hCYP3A4* and creating an indel, Ins8, at the *rCyp3A2* gene. Ins8 proves a convenient tool to demonstrate gene conversion activity for some of the events by molecular means. The high homology between the *rCyp3A1* and *rCyp3A2* allows the gRNA encoded by the AGC cassette to target both loci.

### Configuration of Mouse Active Genetic Target Strains (Figure SM4)

The *mFcgR1* locus had no convenient DNA sequence SNP nearby; so, during the initial construction of the humanized *FCGR1* strain, we created an indel on the chromosome 1.7 kb distance from the mouse *FcgR1* locus by the co-injection with the humanization gRNA a second gRNA targeting a site 1.7 kb distant. Analysis of the humanized animals found several with a successful humanized locus and indels at the 1.7 kb distant site. Among those, we chose a strain with the humanized hFcgR1 minigene and a deletion of 13 bp (or sometimes a 24 bp deletion) at the 1.7 kb site for use during the Active Genetics assessment phase. We also chose to humanize the mouse linked *FcgR2b* and *FcgR3a* loci. Humanizing the linked 83 kb separating *FcgR2b* and *FcgR3*a loci required two sequential rounds of Cas9-assisted substitution, using the single FcgR3a strain as the target for the second round of substitution (Li, et al., 2013). This strategy creates two types of animals, humanized and linked FcgR2b-FcgR3a (cis) and the unlinked-trans configuration (Figure 2).

### F1 strain identity and quality check

All F1 animals in Active Genetics tests were double-checked to show that the genotype planned was the genotype present. This check was conducted for two reasons; first, to assure no strain identity errors were made during breeding steps. Second, our general method to assess Active Genetics (Figure 2) requires the receiver chromosome to have a wild-type sequence and be capable of being hydrolyzed by the gRNA-Cas9 enzyme. All Cas9 strains, especially homozygous strains, might receive a wild-type locus Cas9 induced DSB during F1 strain preparation and before F1 gametogenesis and the F2 breeding test. Thus, all F1 parents with homozygous Cas9, and most of the heterozygous F1 parents, were sequenced at the receiver chromosome target. Only those with wild-type receiver sequences were used for the F1 test cross. Thus, every Cas9 F1 parent was eligible to participate in the gene conversion process.

### Large deletion assay

We observed some genotypes that are inconsistent with usual Mendelian inheritance and Mendelian crossover mechanisms in our F2 test cross. For example, in the F2 genotyping assays for the heterozygous by wild-type cross of rat 3A4-Ins8/WT, we observed genotypes positive for AGC and the downstream marker, Ins8, but negative for both wild-type probes. Since these outcomes are inconsistent with a standard Mendelian model, we investigated the genomic arrangement near the Cas9 target sites.

We hypothesized that a large deletion would eliminate the priming sites for the wild-type probes, see figure SM-1. Taking advantage of the 55 kb between the two Cas9 target sites, we established the large deletion assay (LD) assay. We prepared new PCR primers located +302 upstream of the AGC site and -420 bp downstream of the Ins8 site. We reasoned that if a large deletion occurred, it might be near the target sites and would eliminate the whole 55 kb region; in this case, one could see a short PCR product of about 766 bases, including PCR primers. Indeed, in almost all cases, the short product was present, confirming large deletions presence (SR10). The majority of samples were exactly 766 bps long; a minority were 766 ± 1 to 31 bp. The ∼766 base-pair product was detected as a SYBR green signal in a qPCR assay; then, we confirmed the product’s identity by Sanger sequencing. We set up the special LD assay for the rat *3A1-3A2* AGC. In the mouse, a similar assay was established in strains with AGCs with two gRNA sites in both the cis and trans configuration. No large deletion events were seen in the mouse. Table SR10 summarizes all the cases where Mendelian-inconsistent patterns occurred or might occur in rats and mice, and the assay was conducted to assess the presence of a LD.

### Characterization of Cas9 expression in ovaries and testis

Testis were harvested from wild-type animals, animals with heterozygous or homozygous copies of the Cas9 transgene. Photos (not shown) of testis showed normal size in heterozygous rats, not different from control, for Ddx4-Cas9 heterozygous rats, as-well-as homozygous or heterozygous rats, Sycp1-Cas9 rat, Sycp1-Cas9 mice, and pSycp1-Cas9 bearing mice, and are consistent with the observed fertility. The exception is male homozygous Ddx4-Cas9 rats. Male homozygous Ddx4-Cas9 rats are sterile and have testis about 50% the size of controls. Histology shows no maturing cells beyond spermatogonia in homozygous Ddx4-Cas9 rats (Figure SR5).

### Immunohistochemistry

To assess the expression of Cas9 in tissue, slides were prepared using 10% neutral buffered formalin-fixed tissues that were paraffin-embedded. Five μm sections were used. Antigen retrieval was performed using High PH Leica ER2 (AR9640) for 20 minutes, followed by primary Cas9-directed antibodies for 30 minutes at room temperature and secondary HRP conjugated antibodies. These were then visualized with diaminobenzidine. Immunohistochemistry studies were conducted by Histowiz (New York, New York, USA) using anti-Cas9 antibodies Cat. No. NBP2-36440SS in mouse tissues and AB189380 in rat tissues.

*Cas9 mRNA in tissues* was analyzed from tissues harvested and stored in RNA-Later™. mRNA was prepared using the RNAqueous-4PCR (Thermo Fisher, Waltham, MA, USA) using the manufacturer’s suggested DNAase treatment to suppress genomic contamination. The cDNA synthesis method of the High-Capacity RNA-to-cDNA Kit (ThermoFisher, Waltham, MA, USA) was used. The samples were analyzed using PCR applying primers in the mouse genome on the 5’ side of the Cas9 insert and within the Cas9 sequence. Agarose sizing gels were used to detect and qualitatively assess expression levels.

### Cas9 biochemical activity testis

In testis most of the cells are developing gametes. Hence, we expect to observe DSBs in animals bearing an Active Genetic cassette encoded gRNA and Cas9 expression driven by gamete-specific promoters or drug-regulated promoters. We took advantage of the EJ repair process; we examined testis cells from rats or mice bearing the Cas9 plus an active genetic cassette. In these animals, DNA DSBs at the gRNA site will be repaired by EJ, creating a population of genomic DNA indels. We amplified genomic DNA using PCR primers surrounding the gRNA target site to detect these events. We then used the T7 endonuclease 1 (T7E1) mismatch detection assay (Li, Zhao, et al., 2013; Marshal et al., 1995; reviewed in Sentmanat et al., 2018) to assess activity. Testis single cells suspensions were prepared. Testis were removed and gently burst using the plunger of a tuberculin syringe in 5 mL of RPMI 1640 media and by gentle aspiration in a 5mL serological pipette. The resulting suspension is gently mashed through a 75 um cells strainer and washed with two 5mL aliquots of media, and cells are collected by centrifugation at 350 xg for 5 min. Genomic DNA from testicular cells is prepared using the Direct PCR lysis buffer with proteinase K treatment (Viagen Biotech, Inc., Los Angeles, CA, USA). PCR products were prepared using the testicular genomic DNA and primers from the region around the gRNA target site. They were then assessed with The T7 endonuclease 1 (T7E1) mismatch detection assay, Alt-R Genome Editing Detection Kit (IDT, Coralville, IA), and gel electrophoresis.

### Cas9 expression in testicular cells by DNA content

Testicular single-cell suspensions (∼2 million cells in RPMI-1640 media) were incubated with Vybrant DyeCycle Violet stain (Life Technologies, V35003) for 30 min at 37°C in the dark. Samples were then washed with staining buffer (HBSS + 2% HI-FBS) and fixed in 200µL Cytofix/Cytoperm (BD Biosciences, 554722) for 30 min at RT. Dead cells were removed by counterstaining with Zombie NIR dye (Biolegend, San Diego, CA). Flow cytometric analysis was performed on a Thermo Scientific Attune NxT Flow Cytometer using wavelength settings for channel VL1 for Vybrant DyeCycle Violet, channel BL1 for GFP, and using channel YL for the primary antibody-secondary antibody pair of Cas9 antibody (Novus Biologicals, NBP2-36440SS) and a PE-conjugated secondary (BioLegend, 405307). For gating, we first used FSC-SSC to exclude small debris and large cell clusters, followed by gating for 1N, 2N, and 4N populations based on increasing Vybrant DyeCycle Violet (Rodríguez-Casuriaga, 2014; Bastos et al., 2005; Gaysinskaya et al., 2014). Using the 1N cells as the base, we observe cells we consider 1N, 2N, and 4N to have a per cell MFI ratio (± sd) of 1, 2.11 ± 0.07, and 4.00 ± 0.33 in rats, and 1, 1.97 ± 0.03, 4.02 ± 0.07 in mice. GFP bright cells were captured by setting gates beyond where most events were clustering in no-Cas9 control samples. Data analysis and visualization were performed using FlowJo software (Version 10.7.2, BD).

**Figure SM1:**
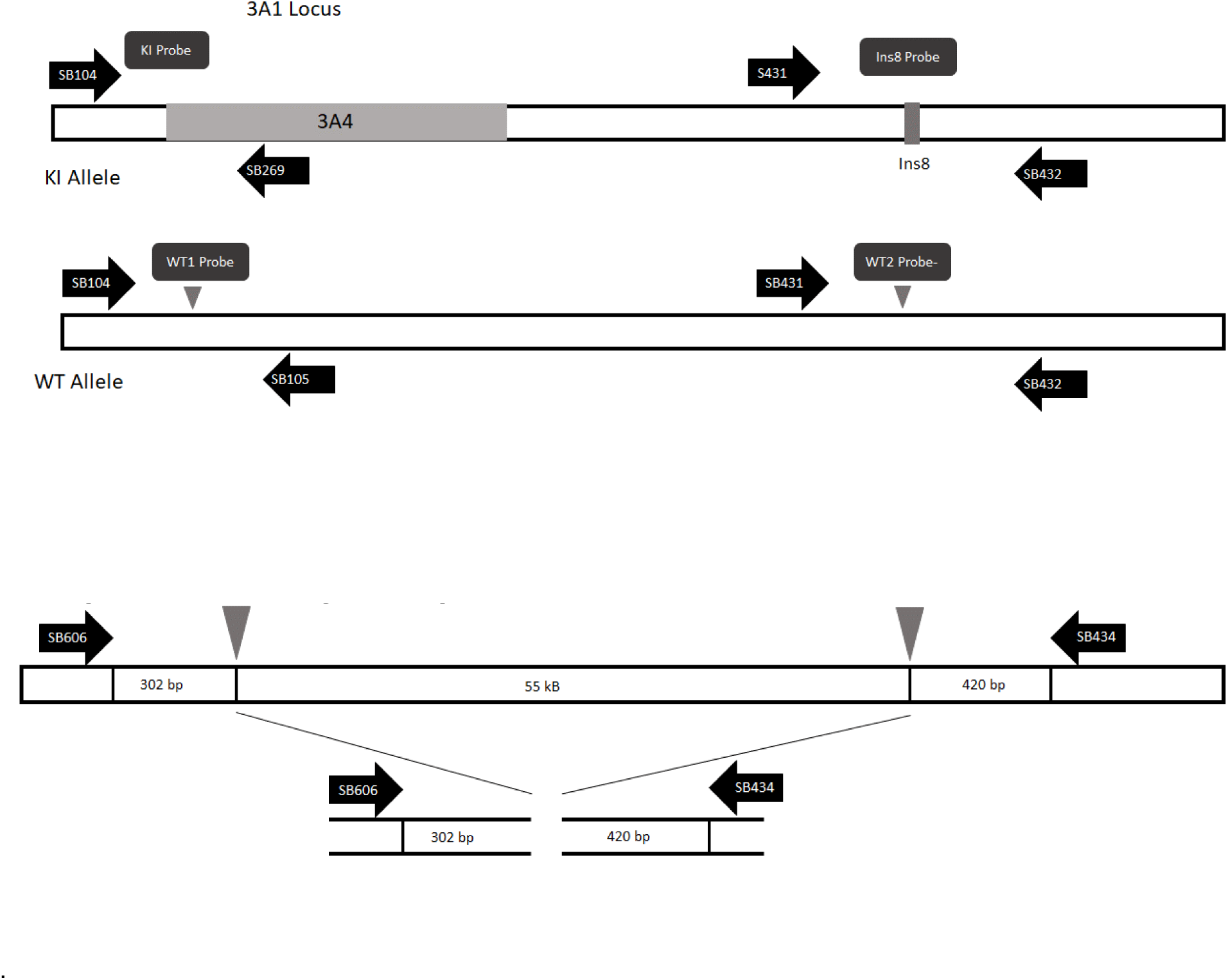
**Panel A.** Arrangements of probes and primer for the standard genotyping assay for Rat *3A1-3A2* locus. Similar genotyping assays were established for each AGC and genetic marker in rats and mice. Samples were scored for the combination of TaqMan^®^ signals assessing the presence or absence of the maker. Figure 1 provides a genotype interpretive guide for cis or trans configured markers. Subsequent Sanger sequencing of the WT locus was conducted for indel detection and further confirmation of the probe findings. Grey triangle, location of Cas9-gRNA target sites. **Panel B: Large Deletion Assay** Arrangement of primers for the Large Deletion Assay for the Rat *3A1-3A2*, detection of a product was scored by forming SYBR green signal, and subsequent Sanger sequencing of the PCR product to characterize the large deletion. Grey triangle, location Cas9-gRNA target sites.

**Figure SM2:**
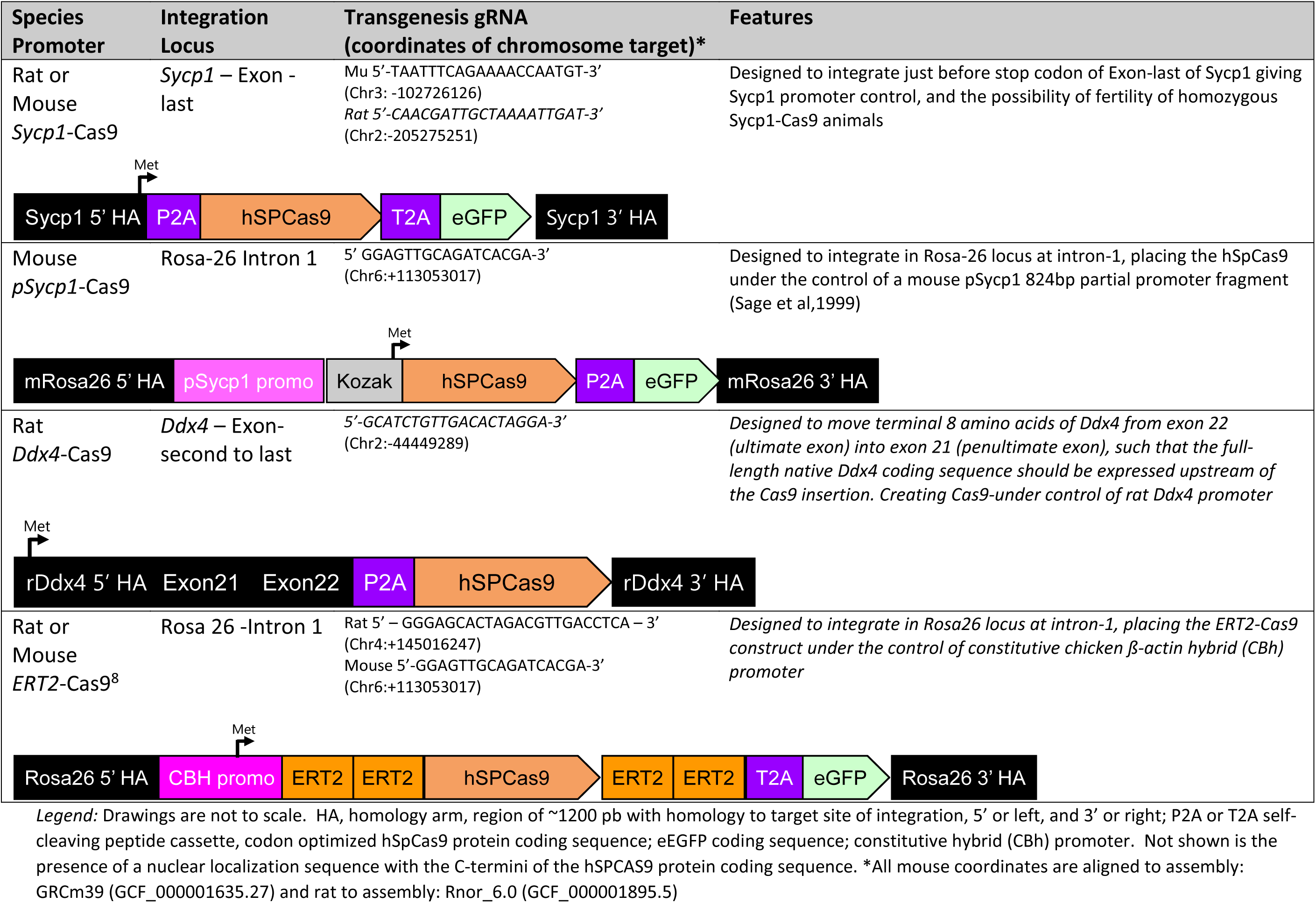
Genetic Arrangement of Cas9 in Rat and Mouse Accelerator strains.

**Figure SM3:**
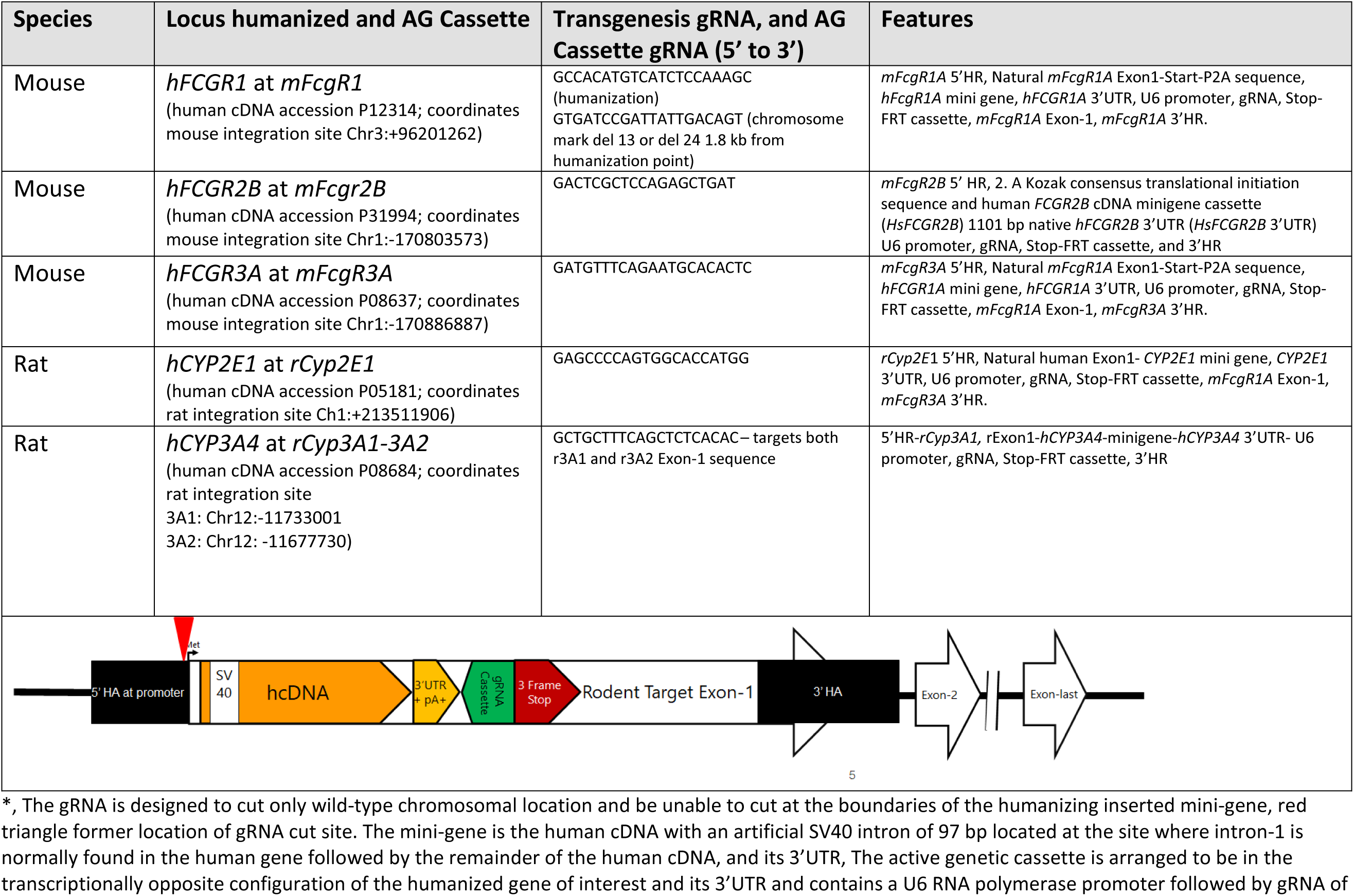
Genetic Arrangement of Humanizing Mini-Genes and Active Genetic Cassettes

**Table SM4:**
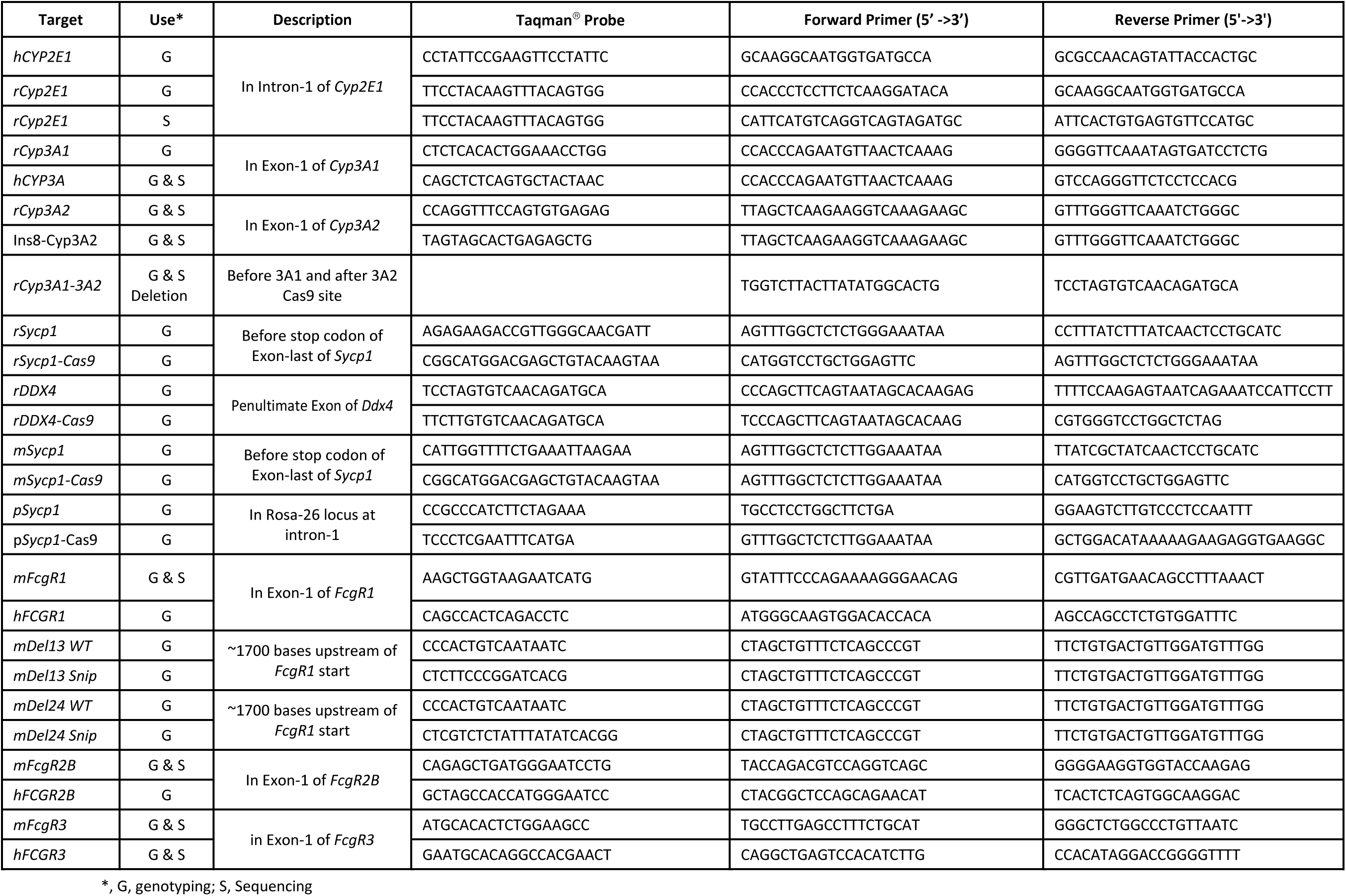
PCR primers and Taqman^®^ probes used in this work

## Supplemental Results

### Fertility

Since many of the Accelerator strains could have changed genes with roles in gamete development, we tracked the fertility of males and females. The Accelerator strains were configured to allow natural expression of the target proteins, SYCP1 or DDX4, using P2A or T2A protein self-cleavage sequences, Figure SM4. The 189 rat parents produced 3265 F2 offspring derived from 341 litters, averaging 9.8 ± 0.3 pups/litter. The heterozygous Ddx4 Cas9 strains bore 10.4 ± 1.2 pups/litter strains of rats with no Cas9 bore 12.5 ± 1.7. This difference is not statistically different, p<0.05. However, the female homozygous Ddx4 strains produced 6.5 ± 1.7 pups/litter; this difference is statistically different at p<0.05. In particular, the homozygous female Ddx4-Cas9 strain bearing the 2E1 AGC produced only 5.3 pups per litter. The Ddx4 litters were 51% male; the male-to-female ratio is normal. In mating cages, they have litters at the same rate as Cas9 absent rats, 0.5-0.6 litters every four weeks. The lower fecundity of Ddx4 females would be a drag on the practical implementation of AG gene conversion using this promoter as arranged in these studies; however, it is offset by the observation of excellent AG efficacy in this strain. Other arrangements of the Ddx4 Cas9 should be investigated. The Sycp1 Cas9 rat strains produced 9.3 ± 1.6 pups/litter; strains of rats with no Cas9 bore 12.4 ± 1.7. This difference is statistically different at p<0.05. The homozygous Sycp1-Cas9 rat strains were fertile but produced about three pups per litter less than non-Cas9 bearing strains. Homozygous *Ddx4* rat males are infertile, exhibit no sperm cells more mature than spermatogonia (figure SR5), and have testis about 50% smaller than wild-type males (not shown).

In mice, the 193 parents produced 3468 pups. The mouse pSycp1 Cs9 lines produced 7.8 ± 1.1 pups/litter, the Sycp1 Cas9 Strain produced 7.7 ± 0.7 pups/litter; strains with no Cas9 produced 7.5 ± 0.7 pups/litter. These differences are not statistically significant, p<0.05. Heterozygous and homozygous strains have similar numbers of pups. These Cas9 strains have no measurable effect on fecundity in mice.

In summary, female rats and mice, homozygous for *Sycp1-Cas9* and *pSycp1-Cas9* strains, are fertile. Homozygous female Rat *Ddx4-Cas9* strains are fertile but produce smaller litters. Male homozygous rat *Ddx4-Cas9* strains are infertile. Mouse heterozygous or homozygous *Sycp1-Cas9* and *pSycp1-Cas9* are fertile.

### Expression assessed by mRNA

To verify our understanding of Cas9 mRNA expression in the organs of rat and mouse Accelerator Strains, we studied mRNA expression in the testis, ovary, and the non-gonadal tissue, the kidney. Cas9 mRNA was strongly expressed in the testis of rats and mice with all four promoters, Figure SR1. Sycp1 homozygous animals expressed more Cas9 mRNA than heterozygous animals in the testis of rat strains. The mouse *Sycp1* homozygous or heterozygous strains express similar amounts of Cas9 mRNA in the testis. In contrast, immune reactive Cas9 is more prominent in the testis of homozygous *Sycp1* strain; the source of this distinction is unclear. The mouse p*Sycp1* driven mRNA was expressed in gonadal tissues and non-gonadal tissues, the kidney. The p*Sycp1* promoter fragment-Cas9 construct was placed in the Rosa-26 locus. This fragment was previously characterized to drive the expression of a reporter in primary spermatocytes cells (Sage et al., 1999). Our histochemical findings show prominent expression in the secondary spermatocytes (Table 2, Figures SR3 and SR5). The source of this difference is unclear; however, the Sage et al. finding was derived from transgenic mice with randomly located genomic integrations, and it characterized only a few strains. In contrast, our method places a single copy only in the Rosa-26 locus; these differences could influence the location of expression. Interestingly, the homozygous Rat *Ddx4-Cas9* strain expresses mRNA for Cas9 even though this strain has no maturing cells beyond spermatogonia and support cells. We detect no testicular immunohistochemical staining for Cas9 in the homozygous cells. The additional sensitivity of mRNA detection vs. immunohistochemical detection may be the source of this difference.

Adult ovaries and oocytes from all strains express some Cas9 as assessed by mRNA analysis and expression detectable by immunocytochemistry in maturing oocytes of the *Ddx4* rat strain and p*Sycp1* mouse strain. Low levels of mRNA expression were seen in adult ovaries of mice bearing Cas9 integrated behind the natural *Sycp1* promoter. However, we can detect no immune reactive Cas9 in mature primary, secondary or antral oocytes (not shown), suggesting carry over or continued expression in mature oocytes is low for the mouse *Sycp1* promoter. Sage et al. (Sage et al., 1999) report that *Sycp1* is expressed in the ovary at E16 and E17, a time during which meiosis is active in primordial oogonial cells.

Cas9 activity and homology-directed repair will promote AG; however, carryover of Cas9 into the developing oocyte, the fertilized oocyte, or the embryo could promote nonproductive EJ events. So, we wished to understand how much Cas9 protein might still be found in late-stage oocytes just before release as mature oocytes. Figure SR4 shows staining of wild-type rat oocytes and *Ddx4*-Cas9 induced expression in the heterozygous and homozygous configuration. Expression in the nearly mature oocyte in the mouse, driven by the *Sycp1* promoter, is not distinguishable from background staining and is not displayed. Cas9 is prominently expressed in the homozygous *Ddx4*-Cas9 oocyte in the cytoplasm and the nucleus. Less but discernible expression is seen in the heterozygous configuration. In the mouse, expression driven by the p*Sycp1*-Cas9 arrangement is detectable in the oocyte cytoplasm of homozygous mice. Expression in the oocyte of heterozygous p*Sycp1*-Cas9 configuration is not distinguishable from wild-type mice. In summary, oocytes within primary follicles or antral follicles of homozygous *Ddx4* rats, heterozygous *Sycp1 rats*, and homozygous *pSycp1* mice express immunoreactive Cas9 (SR4).

**Figure SR1:**
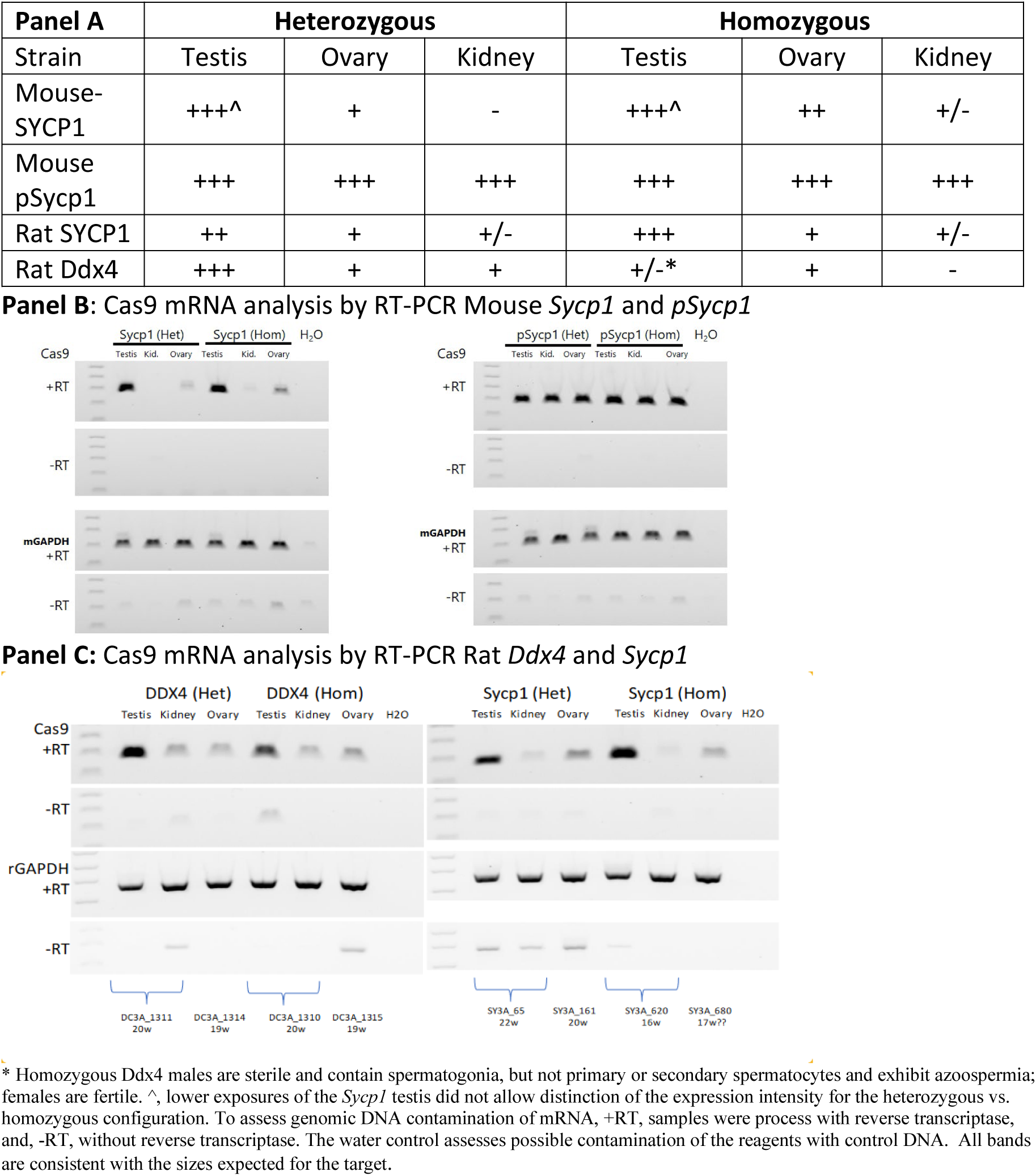
A: Expression of Cas9 mRNA in Accelerator Strains of heterozygous (Het) or homozygous (Hom) rats or mice was collected and is summarized. B: mRNA expression analysis of mouse SYCP1 and pSycp1 accelerator strains, C: mRNA expression analysis of mouse Rat *Sycp1* and *Ddx4* strains.

### Expression of the Cas9 in developing spermatocytes

The cell stage during meiosis I at which Cas9 expression occurs was further characterized by assessing the DNA content of testicular cells and Cas9 or Cas9-GFP surrogate content using flow cytometry. In the rat, *Sycp1* driven expression of Cas9-T2A-GFP expression is significant, showing a mean fluorescent intensity 1.9 times, heterozygous, and 2.9 times, homozygous, the intensity of wild control strains (Table SR6A). In Rat testis from homozygous *Sycp1* accelerator strains, 7.6% of the 4N cells are GFP bright vs. 0.3% in wild-type animals. Substantial fractions of the 1N cells are GFP bright, up to 23% in homozygous animals. 2N cells are 1.8-fold brighter than wild-type animals (Table SR6A). In heterozygous *Ddx4* Cas9 rats, anti-Cas9 immuno-cytochemical detection identifies testicular cells that are 1.8-fold brighter than cells from wild-type animals, indicating substantial expression of Cas9. In Rat testis from heterozygous *Ddx4* accelerator strains, immuno-cytochemical detection of Cas 9 reveals 6.8-fold more of the 4N cells are Cas9 positive versus 4N cells from wild-type animals, 4.8% vs. 0.7% (Table SR6B). Thus, there is expression in leptotene through pachytene stage cells.

In mouse testis from heterozygous *Sycp1* animals, the mean fluorescent intensity is 2.1-fold the intensity observed in wild control animals, 313 vs. 149 (Table SR6A). There are 90-fold more Cas9-T2A-GFP bright 1N cells than in wild control animals, 28% vs. 0.3% (Table SR6A). Seven-fold more 4N cells are Cas9-T2A-GFP bright homozygous strains vs. wild-type 4N cells, 2.7% vs. 0.3%. In the p*Sycp1* mouse, there are 11-fold more 4N cells that are Cas9-T2A-GFP positive compared to wild-type animals, 3.4% vs. 0.3%. Thus, as in the rat, there is expression in leptotene through pachytene stage cells in the mouse.

Readers should note the Cas9 used in our constructs has a C-terminal nuclear-localization sequence (NLS), and in our immune-histological studies, we see nuclear concentrated Cas9. In contrast, GFP protein, liberated upon peptide self-cleavage, is not modified with an NLS; hence, immuno-histological detection shows a widespread distribution of GFP in the cytoplasm and nucleus of testicular cells in *Sycp1* mice, as shown using serial sections in Figure SR7. During post-meiosis I cell maturation of sperm, the cytoplasm is progressively excluded from the cell; thus, the retention and fates of NLS-tagged-Cas9 and GFP may not overlay precisely with the immune-histochemical detection of NLS-tagged-Cas9. Our immunocytochemical detection shows that 4N cells in primary spermatocytes express Cas9-T2A-GFP. The levels of Cas9-T2A-GFP increase in homozygous animals. Our immune-histochemical detection supports these conclusions.

### Biochemical activity of Cas9 in the testicular cells of Rat

In post-puberty testis, most cells are developing gametes. Hence, in animals bearing an Active Genetic cassette and Cas9 expression, we expect to see Cas9-caused DSBs repaired by the EJ process. We examined testis cells from rats or mice bearing the Cas9 plus an active genetic cassette. In these animals, DNA hydrolysis at the gRNA site will be repaired by EJ, creating a population of genomic DNA indels. We amplified genomic DNA using PCR primers surrounding the gRNA target site to detect these events. We then used the T7 endonuclease 1 (T7E1) mismatch detection assay to assess Cas9 activity. (Li, et al., 2013; Marshal et al., 1995; Sentmanat, et al., 2018). Figure SR8 shows findings from *Ddx4*-Cas9-h2E1 rats and *Sycp1*-*Cas9*-*hCYP3A4* strains. Testicular DNA contains EJ repaired indels at the location of the gRNA target site in the rat *Ddx4*-Cas9 and *Sycp1*-Cas9 strains, but not in strains where the gRNA was absent. Together with our examination of the presence or absence of indels at the gRNA target site in F2 AG test animals, these findings provide evidence that gRNA-Cas9 was present and biochemically active in developing rat sperm (Table SR9A, 9B).

### Large and Very Large Deletions

Large and putative very large deletions occurred only in the cis 3A1/3A2 targeted rat setting. No mouse configuration, either cis or trans arranged, exhibited large or putative very large deletions, Table SR10. We observed a few pups with a Mendelian-inconsistent genotype in heterozygous and homozygous male or female rats with the *Ddx4*-Cas9 or *Sycp1*-Cas9 operating with the *CYP3A4* ACG, targeting the cis r3A1/3A2 locus and. In the heterozygous and homozygous female or male F1 parents expressing *Ddx4*-Cas9 operating with the *CYP3A4* ACG, there were 40 cases observed out of 955 F2 pups examined, Table SR9. In the *Sycp1* driven Cas9 setting acting with the *CYP3A4* AGC, we observed 22 out of 916 F2 pups studied. The large deletion LD products were sequenced to confirm that in the product formed, the -302 and +420 bp primers were now close enough together to allow a product across what was formerly a 55 kb region. Most sequences show direct abutment of the two gRNA target sites, and a minority show inserts or deletions of 2 to ∼30 bases, as expected. Of course, this assay has limitations; there could be deletions even larger than 55 kb; we saw 16 events where no PCR product was formed. We consider these to be putative very-large deletion products (VLD). Thus, in the situation where the rat CYP3A4 AGC was the Cas9 target, the cis r3A1/3A2 locus, 79 deletion cases were observed (62 LD and 17 VLD) compared to 315 Mendel imputed plus molecular Active Genetic events in the 1871 F2 pups. We saw zero such events out of 216 rats in the absence of Cas9 in strains targeting the r3A1/3A2 locus, so the gRNA-Cas9 hydrolysis activity is necessary to form these products. We note that gene conversion-AG events are much more common than these large and very large deletions. These special cases occur at a 25.1% rate as compared to AG.

### Other considerations

During these investigations, we considered other elements of the HDR and crossover recombination pathways that might influence the success of various configurations; recombination hotspots and distances to chromosomal ends might contribute to efficiency. In the Sprague-Dawley rat, no hotspot maps are available; strain-to-strain differences are very large, so data from other strains are not applicable (Mihola et al., 2021). We examined the mouse recombination hotspot map from the c57BL derivative mouse line Hop2 -/- (Smagulova et al., 2011). In the mouse, the sites we tested are 26 Mb to 201 bases from our gRNA hydrolysis sites, *mFcgR2b* is 26 Mb, *mFcgR3a* is 11 Mb, and *mFcgR1* is 221 bases distant. AG gene conversion efficiency is exhibited by the *FcgR2b/3a* combination in some configurations, but the Cas9 hydrolysis sites are 11-26 Mb distant to a hotspot. *FcgR1* showed no AG gene conversion by excess analysis and only three molecularly detected events **but** it is located 221 bases from a hotspot. So, closeness to a recombination hotspot may be only a minor contributor to AG gene conversion efficacy.

A crossover must happen to terminate a BIR event, or the chromosome ends must be reached. We considered the location of our gene conversion targets relative to the ends of chromosomes. In the mouse, m*FcgR2b* is 24.25 Mb, *FcgR3*a is 24.17 Mb, and *mFcgR1* is 63.4 Mb from their chromosomal ends. In the rat, *Cyp2E1* is 68.2 Mb, *Cyp3A1* is 11.73 Mb, and *Cyp3A2* is 11.68 Mb from their chromosomal ends. These distances are quite long; thus, our configurations may not have thoroughly tested the influence of distance to chromosomal ends.

### Considerations related to the practical application of this technology

During our work, as described in the methods, we were vigilant to exclude F1 parents that contained a Cas9-gRNA-derived cut and then an EJ-derived indel on the receiver chromosome. Such indels are blocked from participating in the AG gene conversion process and thus could be a drag on the practical application of the AG technology. We observed between 8% and 40% of the F1 rat parents with a receiver chromosome indel (Table SM11). In mice, we observed between 0% and 63% of the F1 parents had a receiver chromosome indel (Table SR11). In a practical application of the AG gene conversion process wherein one assembles many loci into a single strain, individual animals will need to be tested before making crosses. These kinds of receiver chromosome indels would block the AG gene conversion process. Establishing a practically useful system will have to carefully consider an optimum way to bring the Cas9 gene into the strains bearing the AGCs to avoid unplanned indels, which will impede the overall efficacy of the process. Also, as a practical matter, it is nearly imperative to have the Cas9 on a different chromosome than any of the AGC(s). Making chromosomally colocalized AGC-Cas9 pairs requires either fortuitous crossovers during breeding or laborious sequential assembly. Our FcgR1-Del13 and Sycp1-Cas9 mouse is an example where fortuitous crossover allowed the assembly of this mouse chromosome-1 linked pair.

### AG gene conversion efficacy and parents

Since a reproducible technology is an important goal. Reproducibility would be reduced by high individual-to-individual variability; it could also point to unexpected properties. Thus, we examined AG gene conversion efficacy arising from each parent (Table SR11). For the Cas9 strain AGC combinations in which we studied five or more Cas9-bearing parents, we assessed the distribution of AG efficiencies and looked for skewed distributions. In particular, we looked for parents with consistent 0% AG efficiency or 100% AG efficacy; both cases could indicate system anomalies. The maximum and minimum efficiency data for Cas9-AGC combinations with five or more parents showed litters with higher and lower AG efficiency and were consistent with the mean (Table SR11). We did not observe any Accelerator-AGC combinations with individual parents that exhibited only 0% or 100% AG efficiency, suggesting that odd performance and unexpected strain-to-strain variances are not frequent

**Figure SR2:**
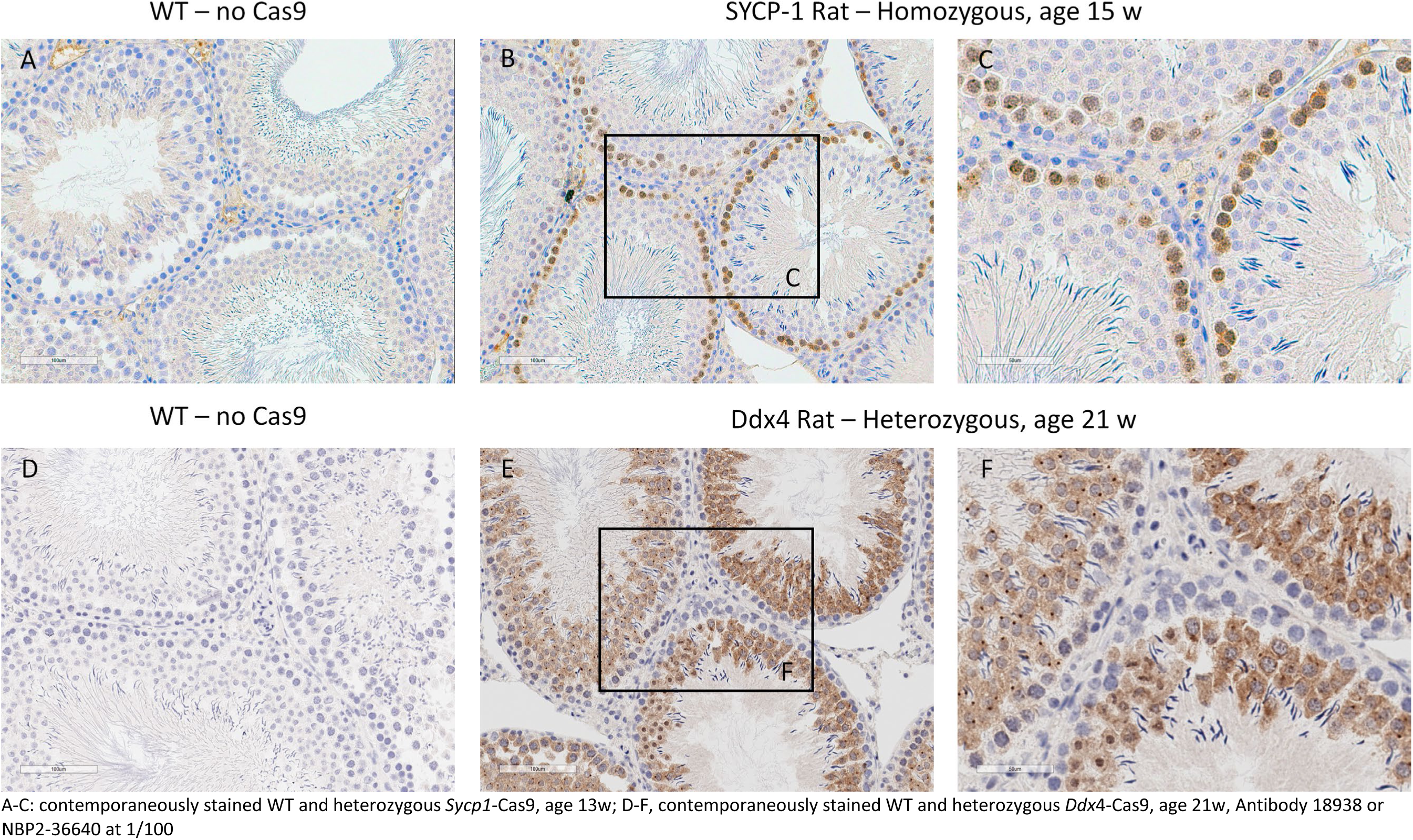
Expression of Cas9 in the testis of **rat** accelerator strains *Ddx4-*Cas9 and *Sycp1*-Cas9

**Figure SR3:**
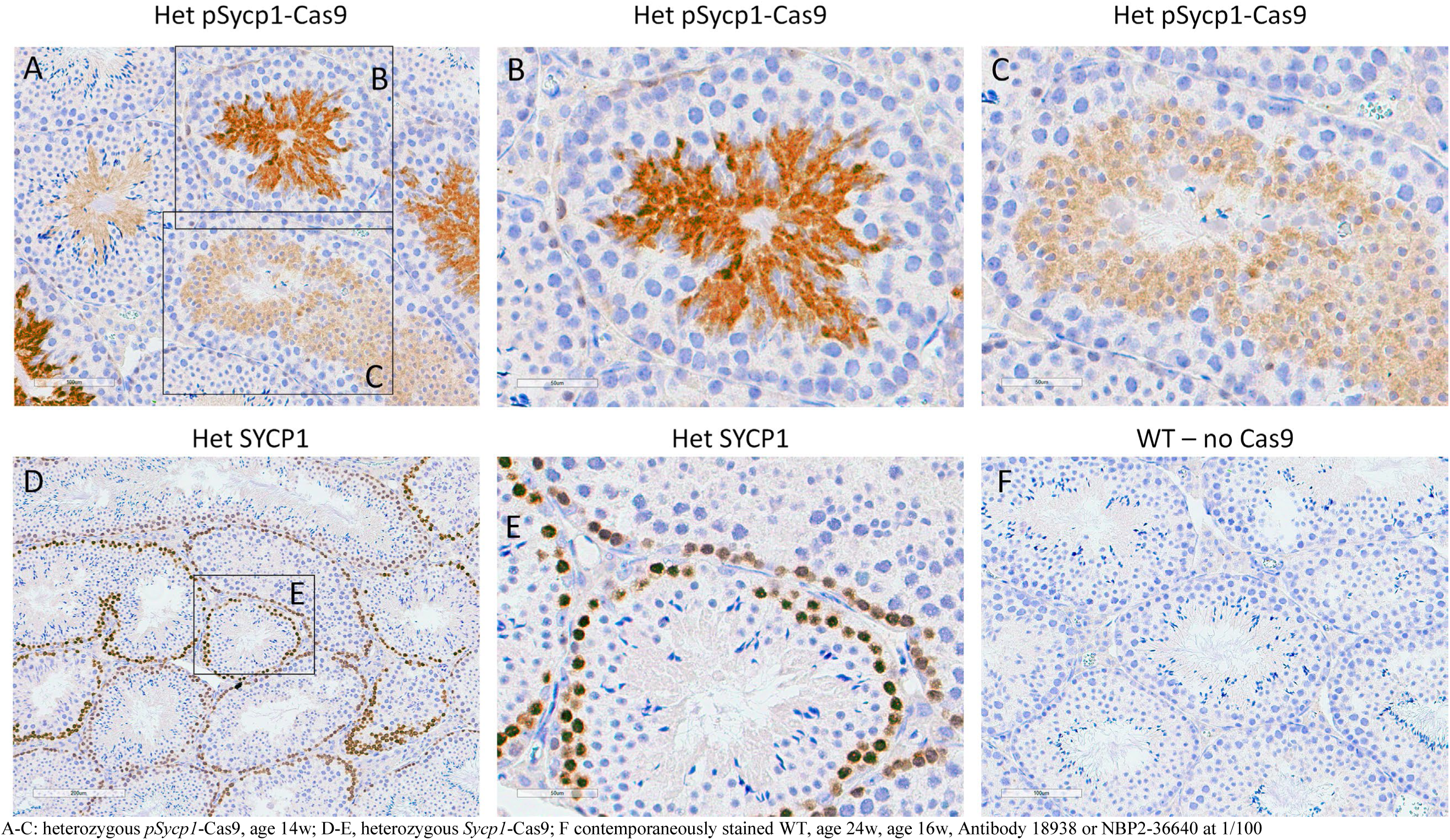
Expression of Cas9 in the testis of **<u>mouse</u>** accelerator strains pSycp1-Cas9 and SYCP1-Cas9

**Figure SR4:**
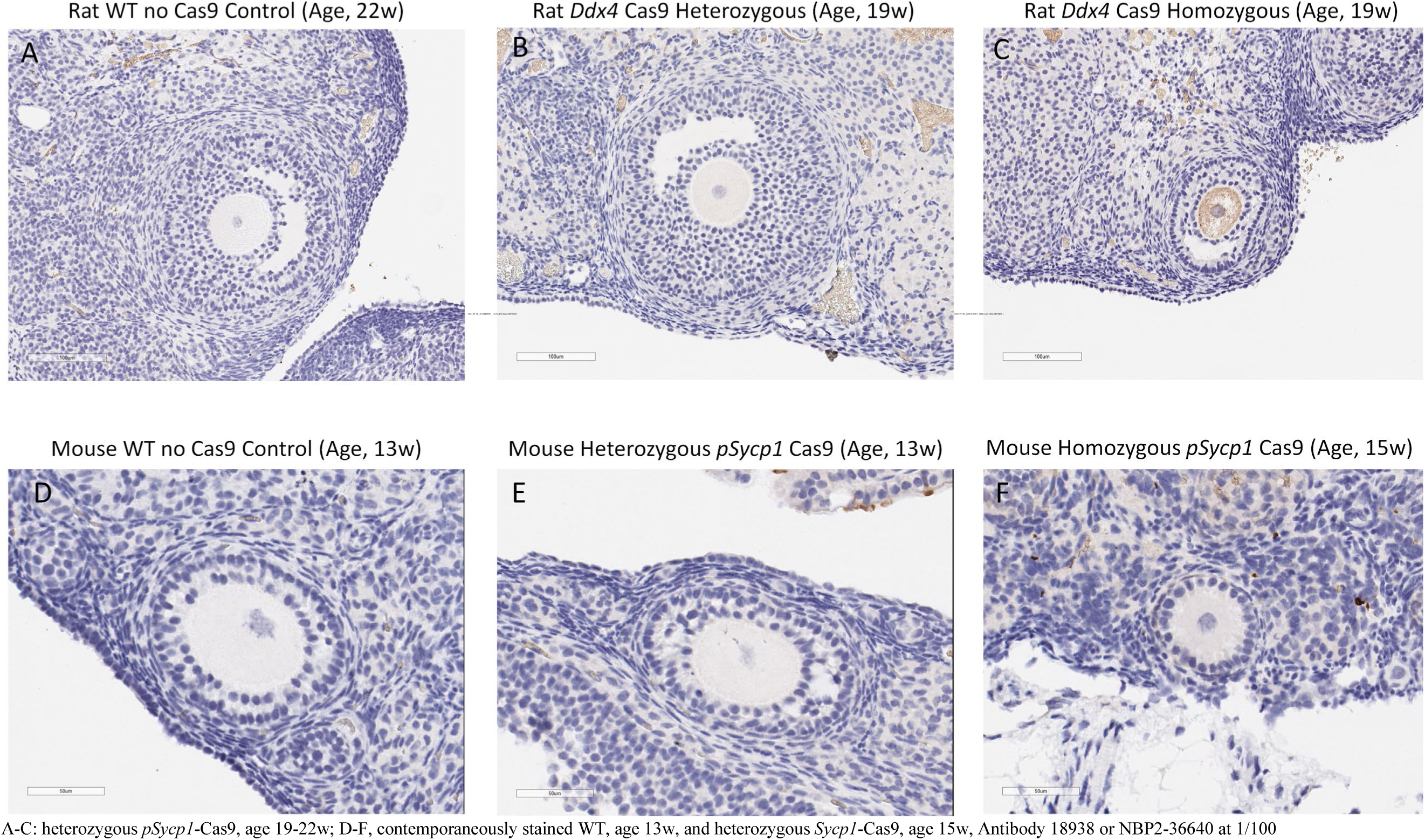
Expression of Cas9 in the maturing oocytes of rat and mouse accelerator strains *Ddx4*-Cas9 and *pSycp1*-Cas9

**Figure SR5:**
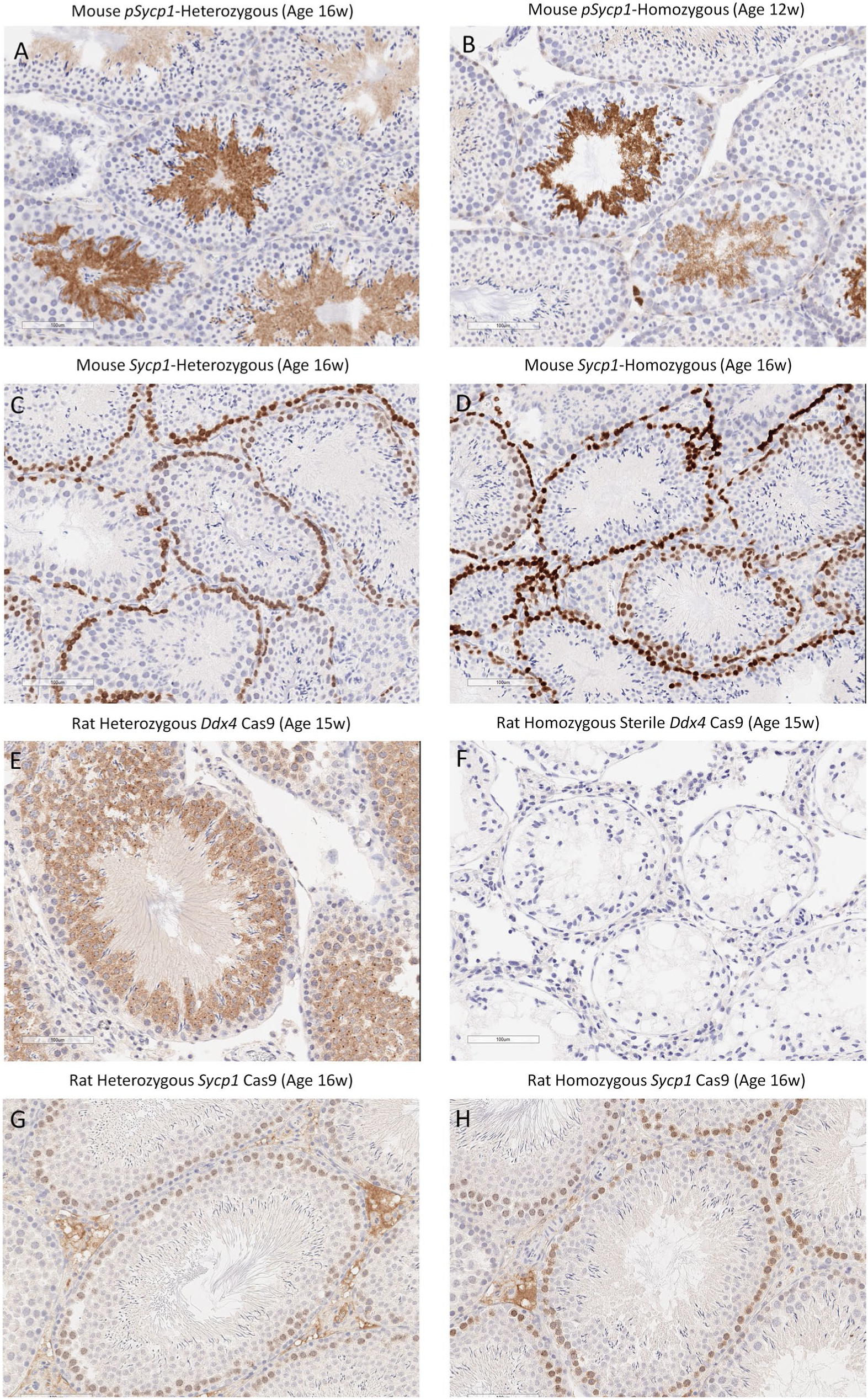
Heterozygous versus Homozygous Cas9 immunohistochemistry in Mouse and Rat (All pairs stained contemporaneously)

**Table SR6A:** **Rat Analysis** of Cas9 co-cistronic green fluorescent protein (GFP), in testicular cells sorted according to DNA content.

| Species | Promoter & Zygosity | GFP MFI entire population | Percentage of 1N cells that are GFP bright (GFP MFI of all 1N cells) | Percentage of 2N cells that are GFP bright (GFP MFI of all 2N cells) | Percentage of 4N cells that are GFP bright (GFP MFI of all 4N cells) |
| --- | --- | --- | --- | --- | --- |
| Rat | WT | 195 | 1.12 (217) | 0.21 (144) | 0.31 (289) |
|  | SYCP1 Het | 369 | 11.1 (500) | 2.58 (216) | 2.73 (383) |
|  | SYCP1 Hom | 568 | 23.1 (928) | 4.94 (263) | 7.45 (434) |
| Mouse | WT | 149 | 0.31 (161) | 0.20 (105) | 0.31 (202) |
|  | SYCP1 Het | 230 | 20.1 (340) | 1.69 (122) | 1.30 (211) |
|  | SYCP1 Hom | 313 | 28.0 (523) | 2.83 (145) | 2.16 (212) |
|  | pSYCP1 Het | 223 | 7.24 (258) | 3.03 (157) | 2.72 (243) |
|  | pSYCP1 Hom | 213 | 7.81 (252) | 3.37 (153) | 3.41 (235) |

MFI, Mean Fluorescent Intensity; Rats were between 16 weeks of age. Mice were between 14 and 16 weeks of age.

**Table SR6B:** **Rat Analysis** of Cas9 in Ddx4 heterozygous animals by Immunocytochemical analysis using anti-Cas9 antibodies, in testicular cells sorted according to DNA content.

| Species | Promoter & Zygosity | Cas-9 Signal MFI entire population | Percentage of 1N cells that are bright in Cas-9 signal (MFI of all 1N cells) | Percentage of 2N cells that are bright in Cas-9 signal (MFI of all 2N cells) | Percentage of 4N cells that are bright in Cas-9 signal (MFI of all 4N cells) |
| --- | --- | --- | --- | --- | --- |
| Rat | WT | 321 | 0.88 (367) | 1.01 (212) | 0.73 (408) |
|  | Ddx4 Het | 586 | 21.5 (668) | 7.00 (352) | 4.81 (542) |

MFI, Mean Fluorescent Intensity; Rats were 18 weeks of age.

**Figure SR6C:**
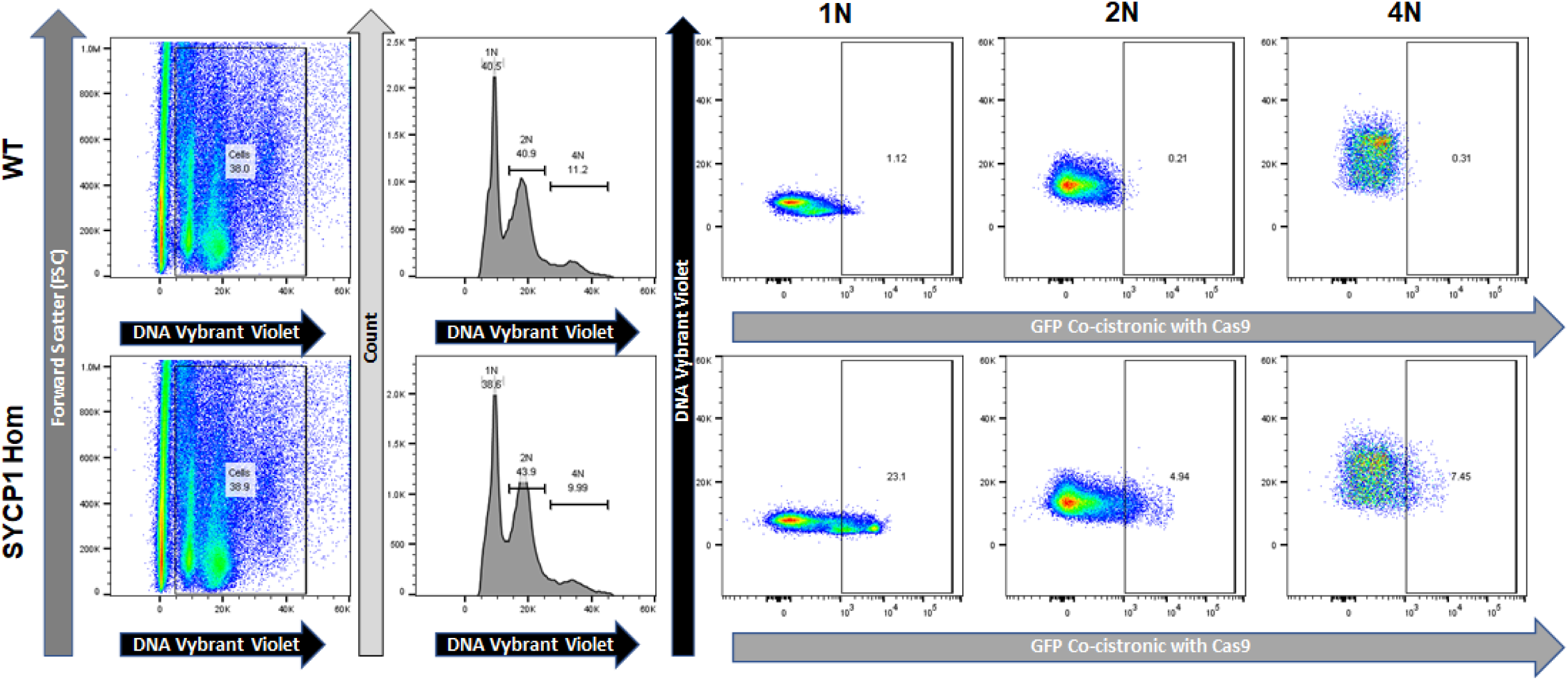
Example rat studies measuring Cas9 co-cistronic green fluorescent protein (GFP), in testicular cells sorted according to DNA content.

**Figure SR6D:**
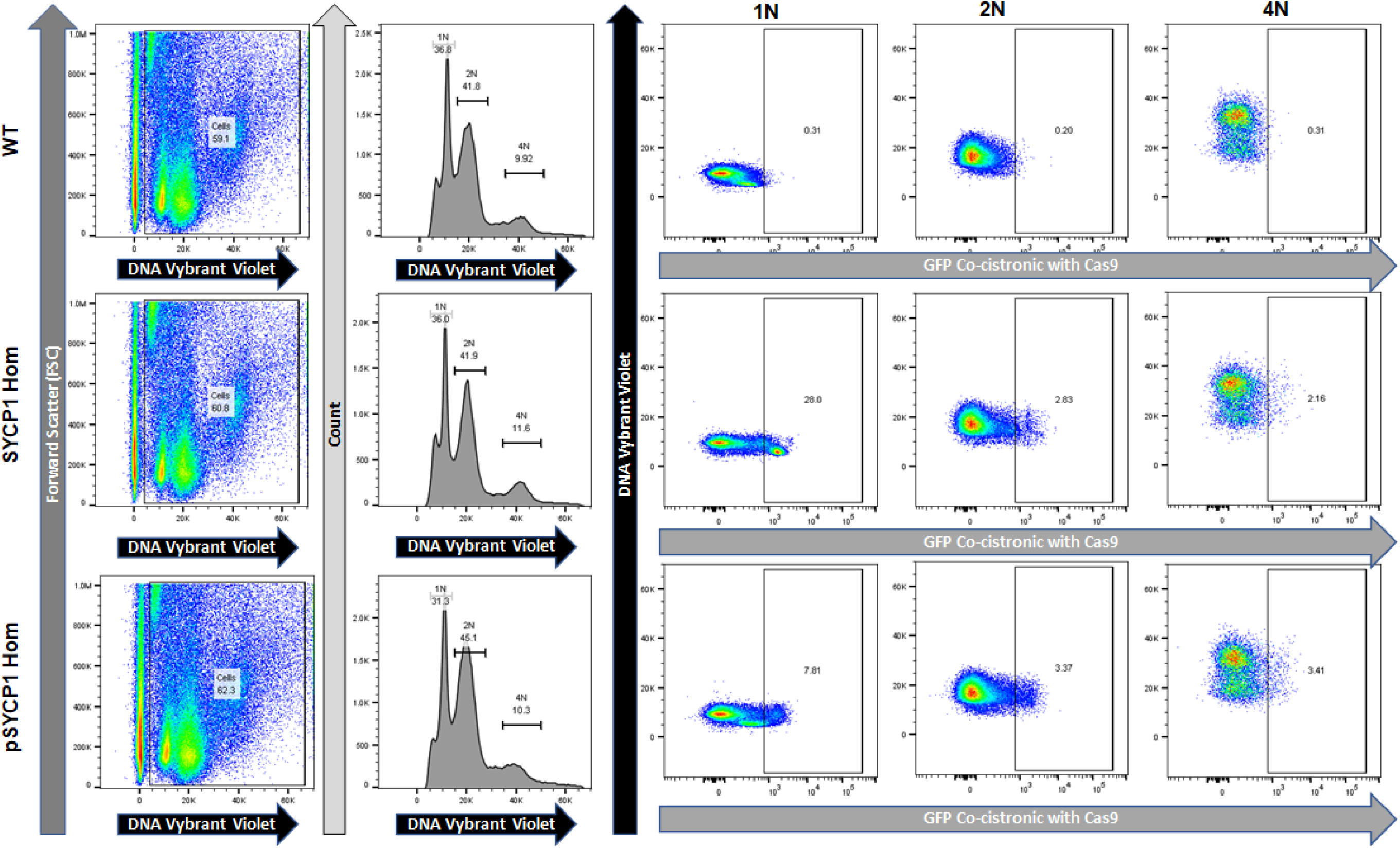
Example mouse studies measuring Cas9 co-cistronic green fluorescent protein (GFP), in testicular cells sorted according to DNA content.

**Figure SR6E:**
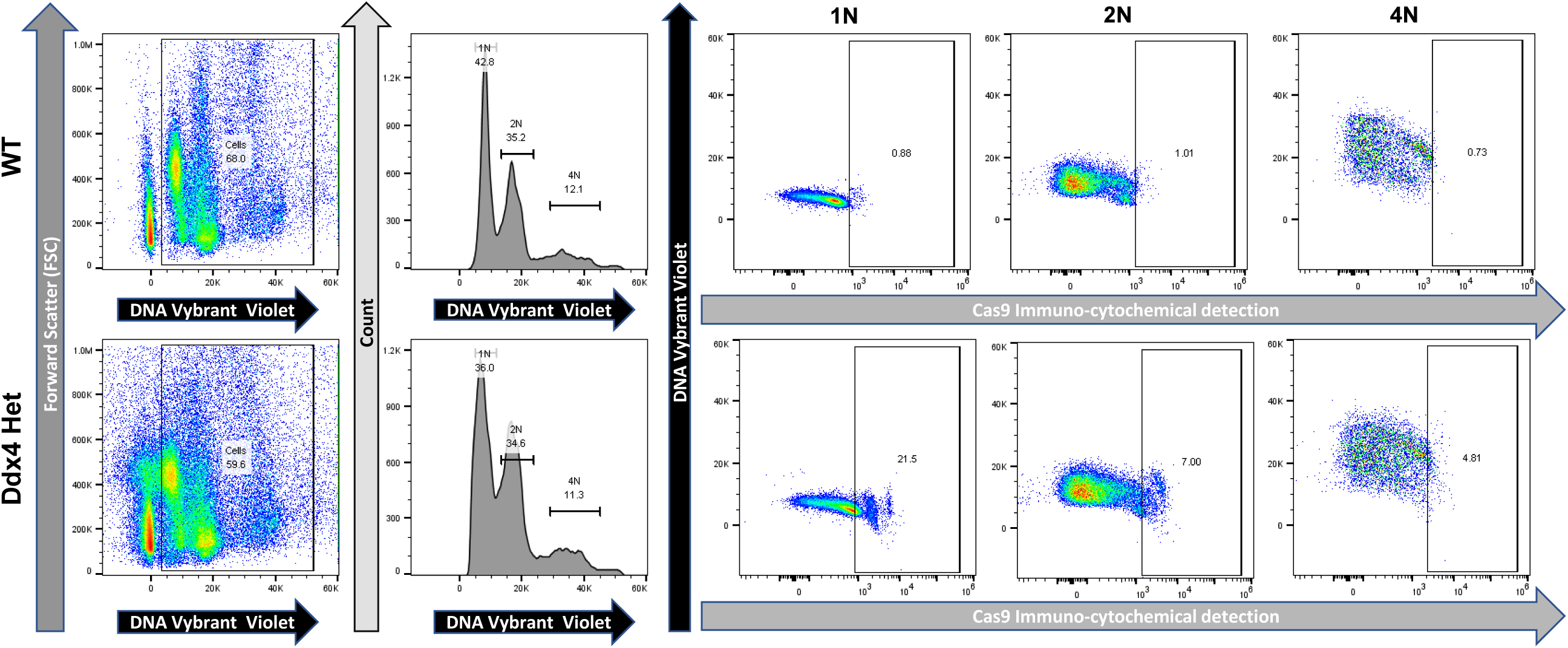
Rat studies measuring Cas9 protein detected immunocytologically in testicular cells sorted according to DNA content.

**Figure SR7:**
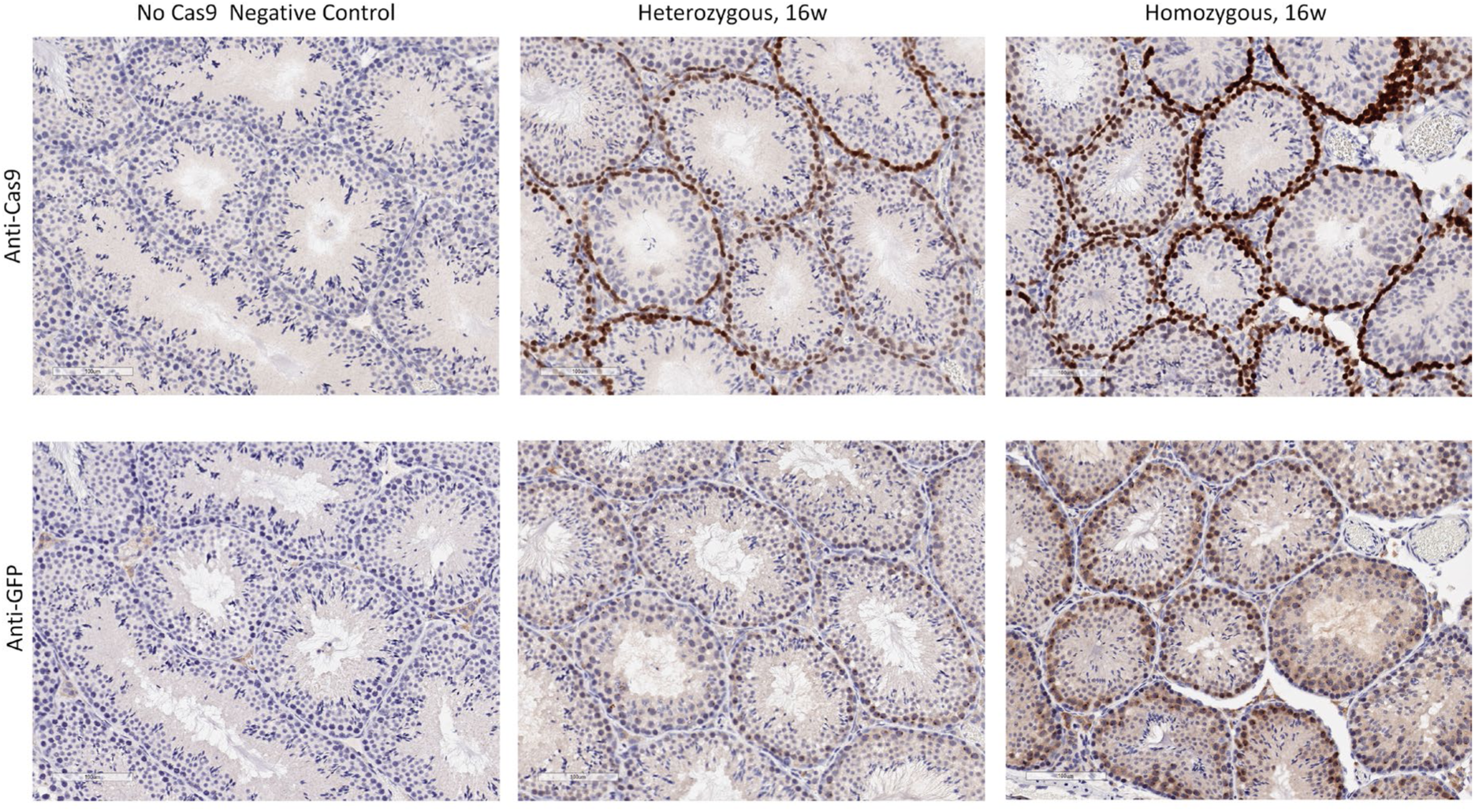
Immuno-histochemistry examination of the expression of Cas9 and co-cistronic GFP in mouse *Sycp1*-Cas9 strains. **Serial sections** were stained with Anti-Cas9 (upper row) or Anti-GFP antibodies (lower row).

**Figure SR8:**
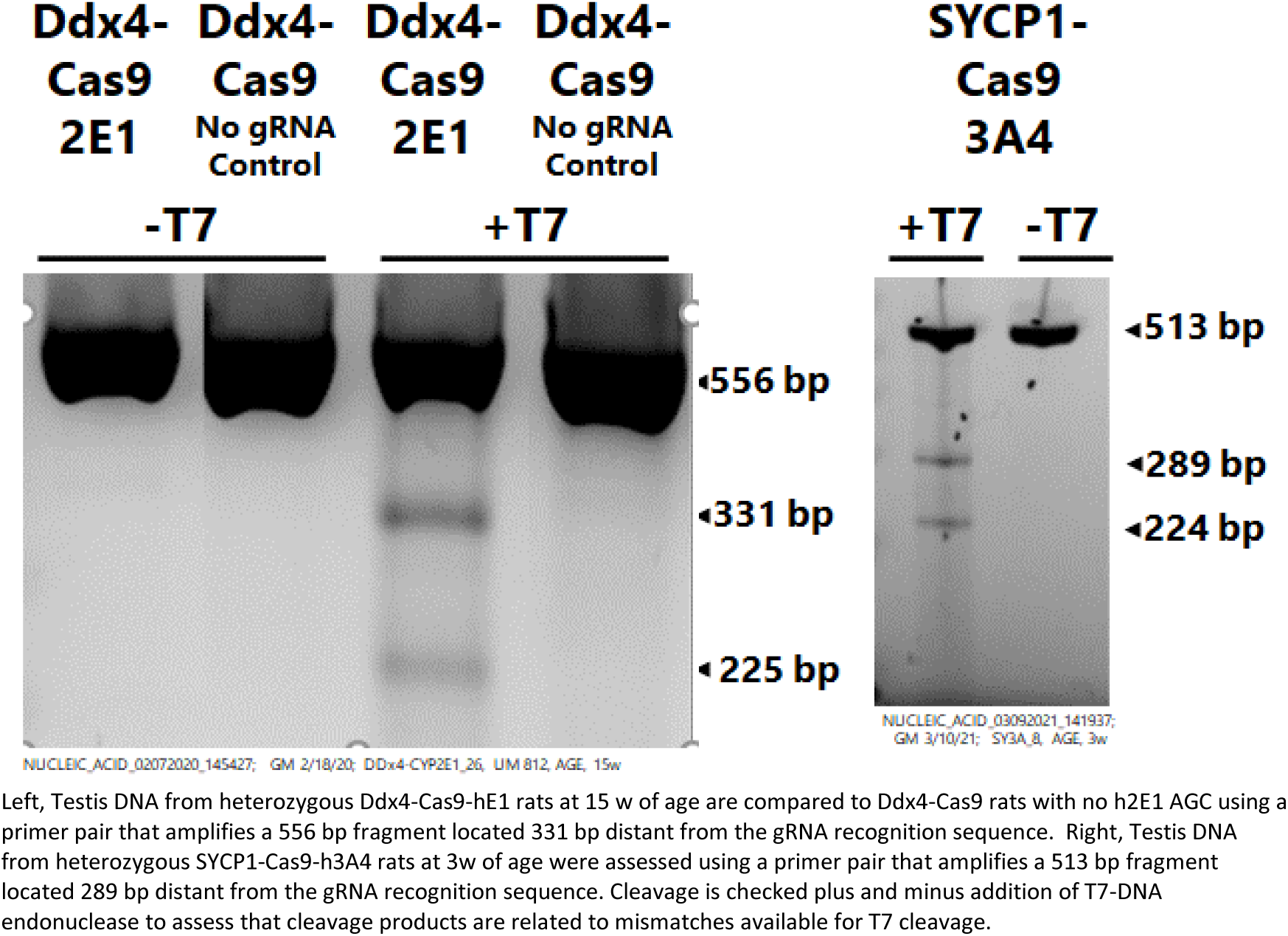
Detection of biochemically active gRNA-Cas9 in the testis of rat strains bearing Ddx4-Cas9-2E1, Ddx4-Cas9-no AGC, or SYCP1-Cas9-3A4 strains.

**Table SR9A:**
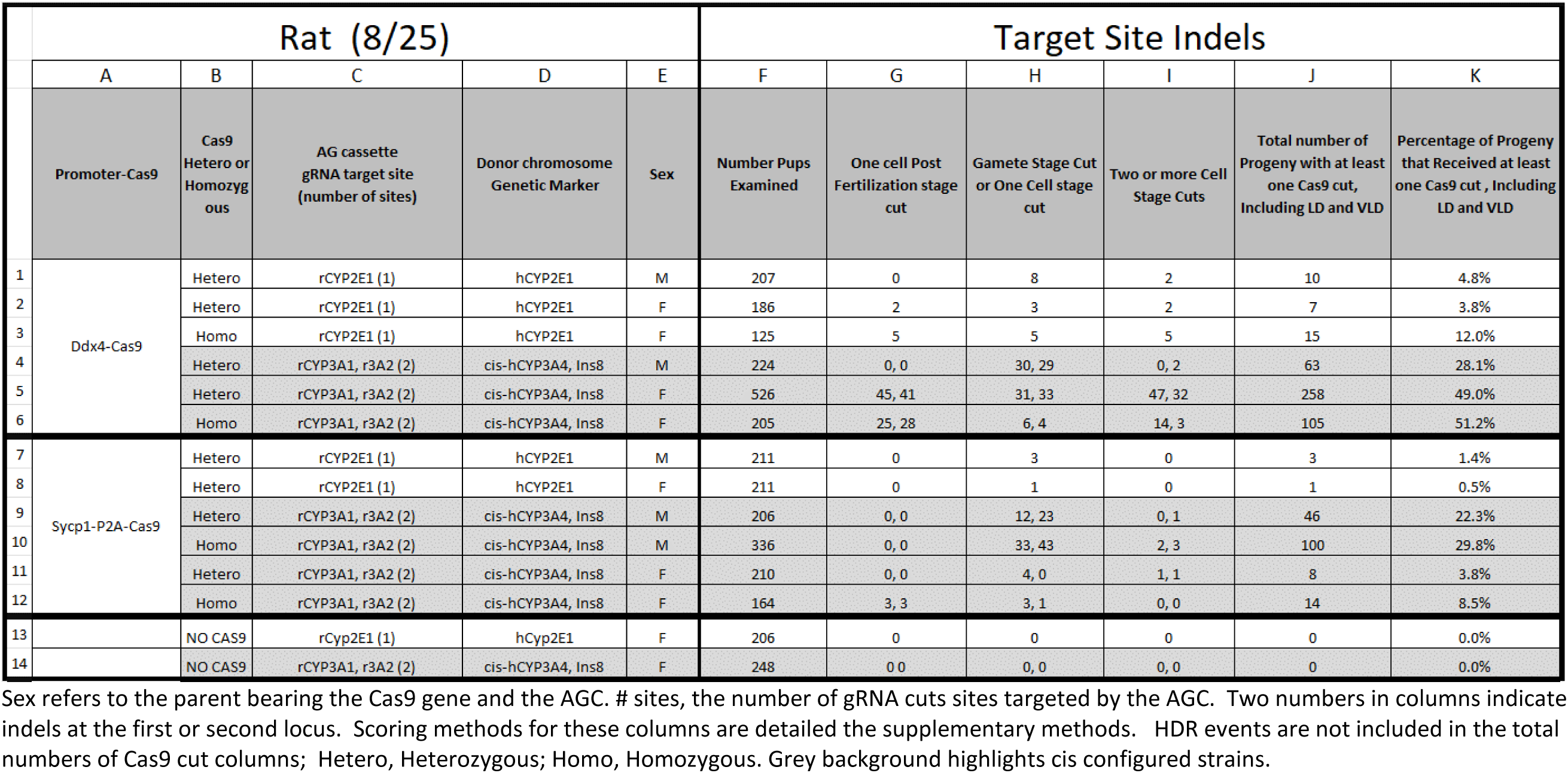
**Rat Indels** -- Influence of Cas9 promoter, Cas9 gene dosage, locus, and parent sex on the occurrence of Cas9 induced indels, and development stage at which Cas9 activity was present.

**Table SR9B:**
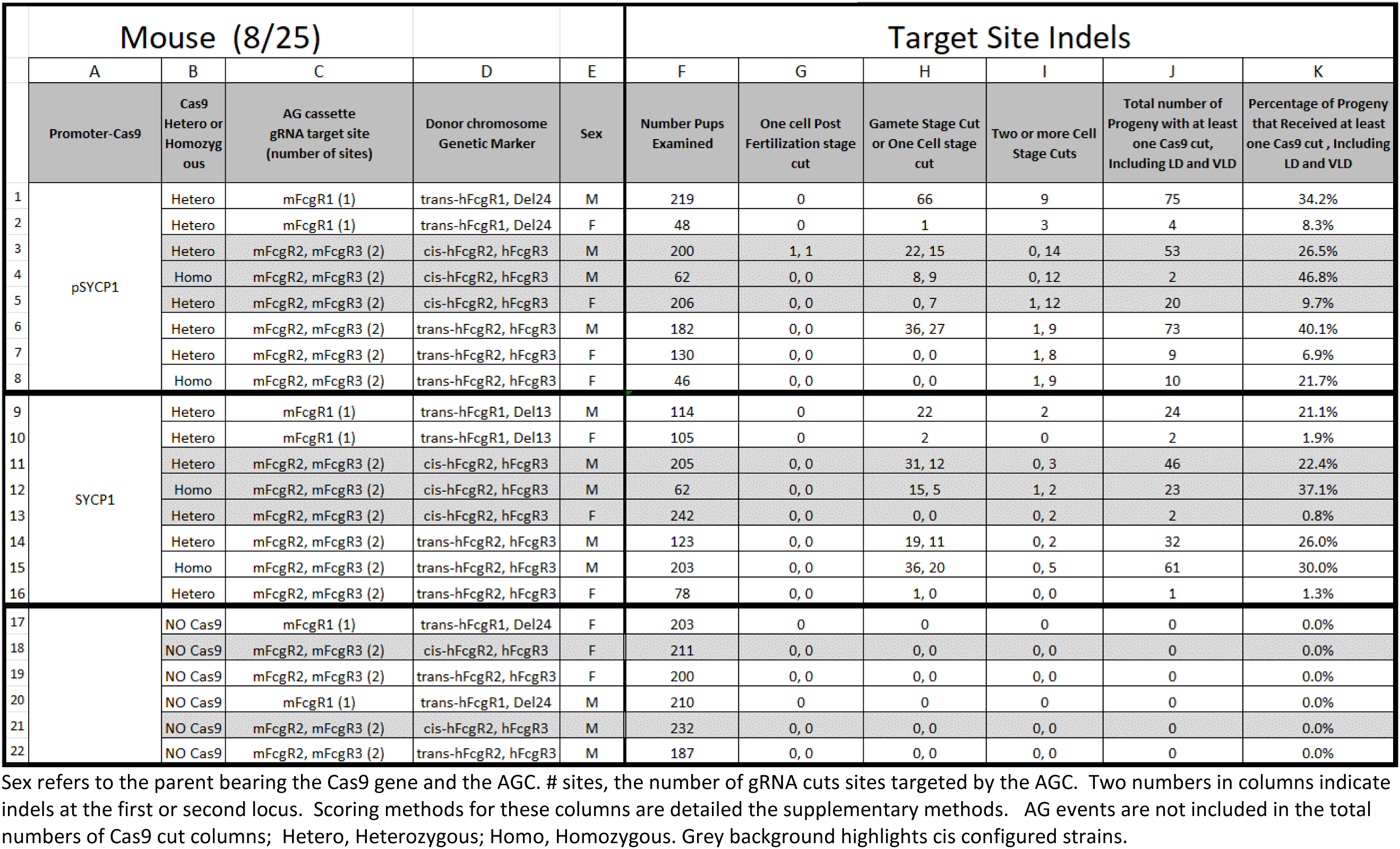
**Mouse Indels** -- Influence of Cas9 promoter, Cas9 gene dosage, locus, and parent sex on the occurrence of Cas9 induced indels, and development stage at which Cas9 activity was present.

**Table SR10.**
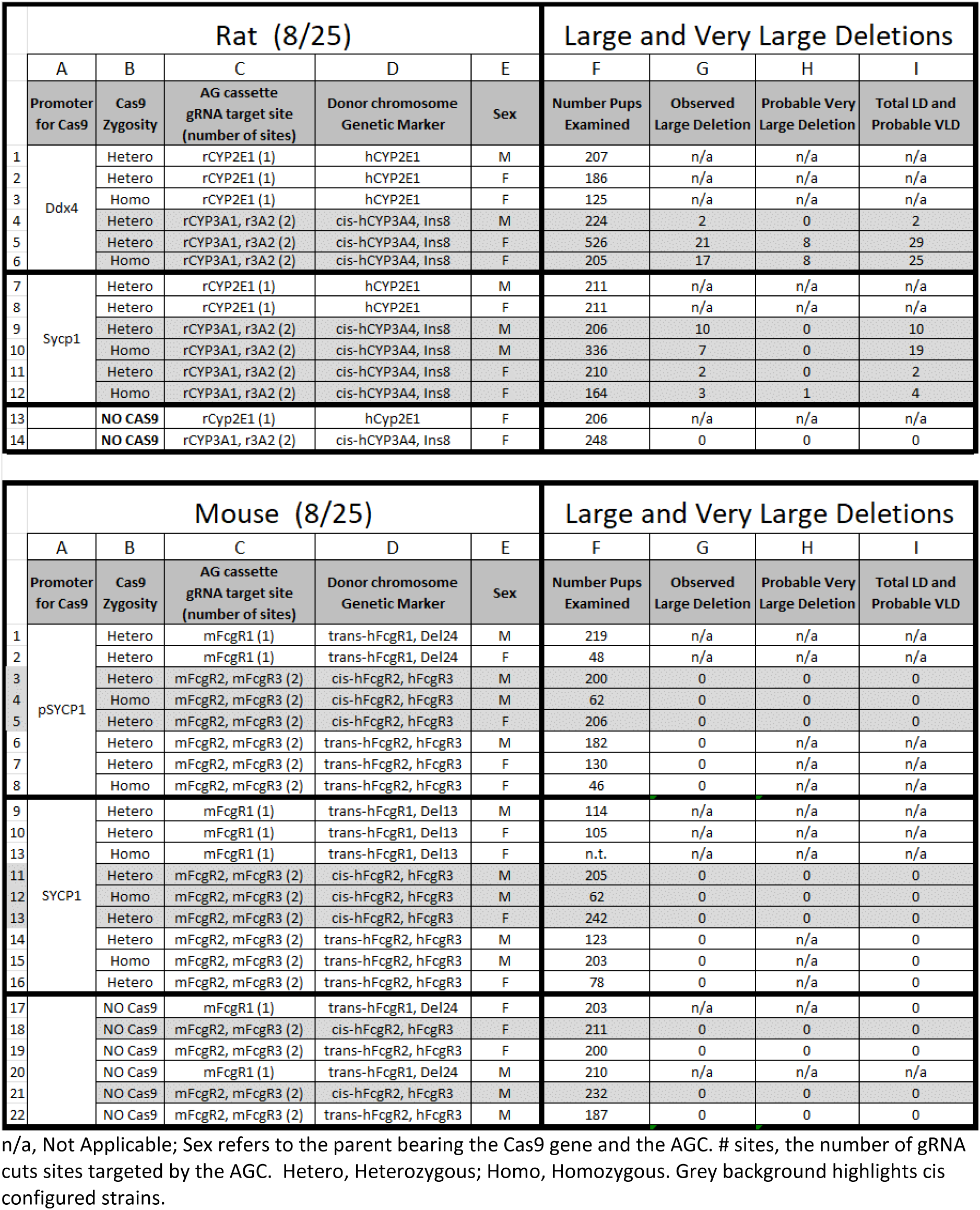
Presence of Large and Very-Large Deletion in strains that vary Cas9 promoter, Cas9 gene dosage, locus, parent sex, and species

**Table SR11:**
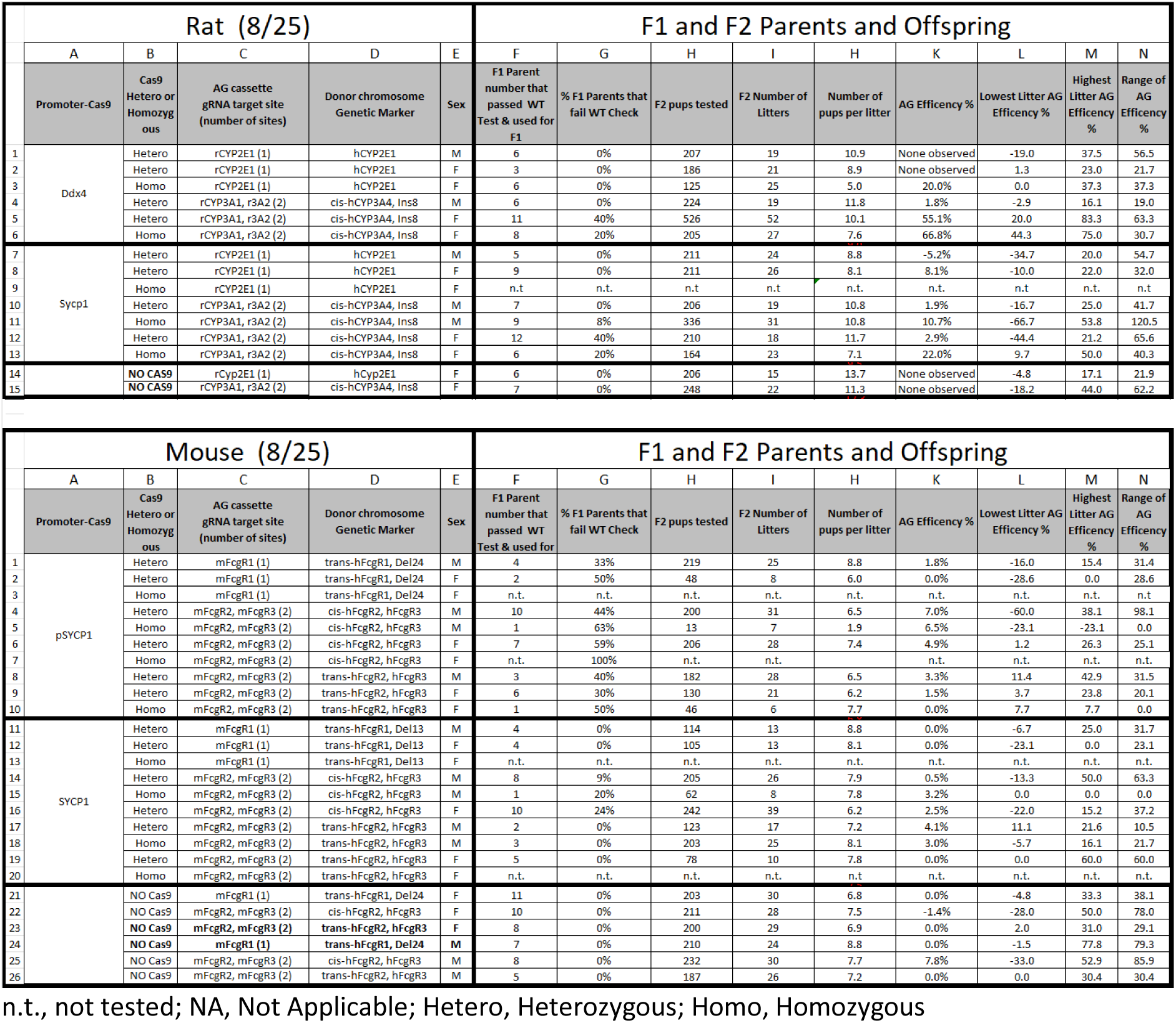
Characterization of F1 parents, F2 litters, and AG efficiency by parent and litter basis

**Figure SR12:**
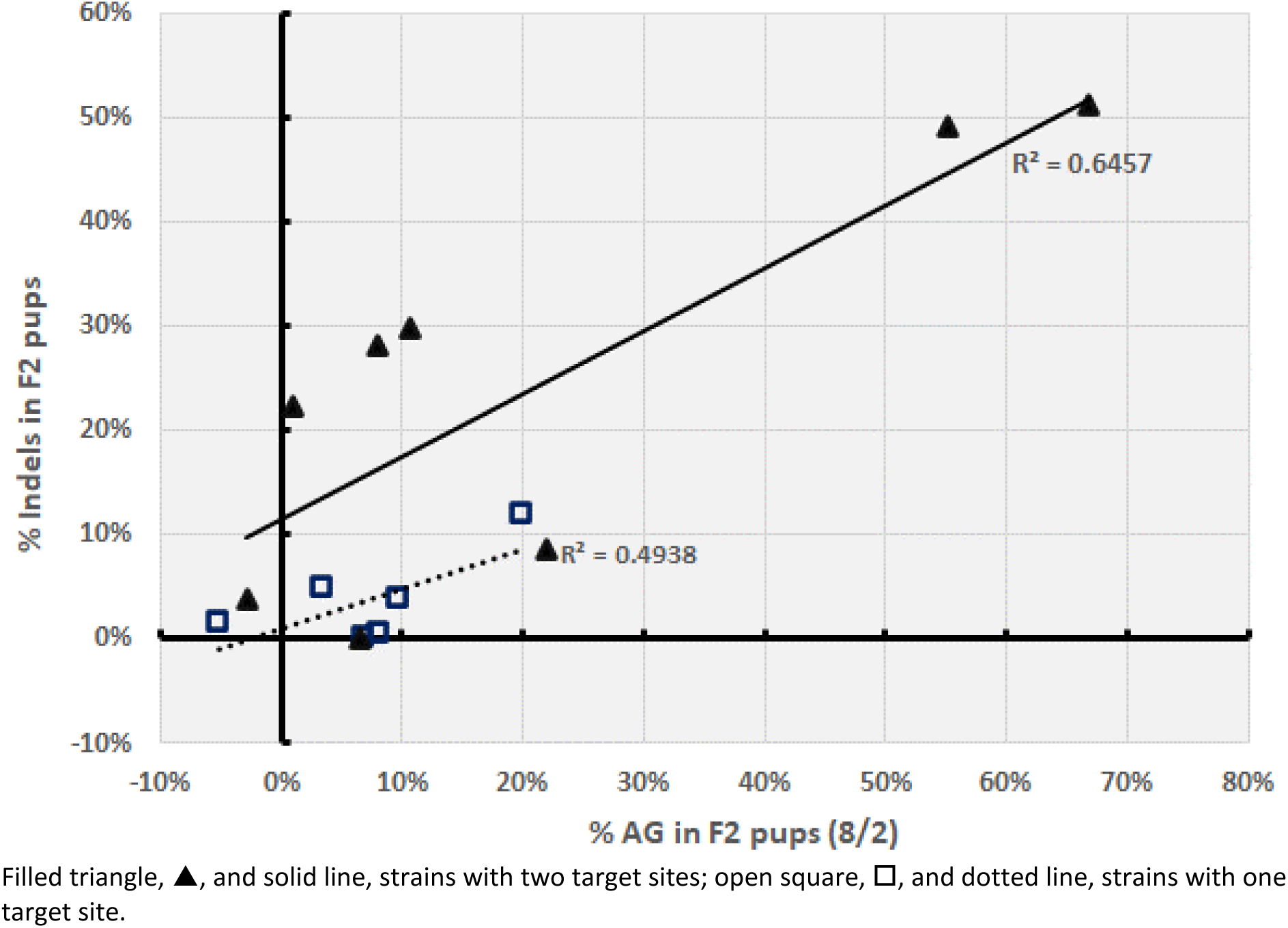
Correlation between Cas9 Derived Indels and Gene Conversion in the Rat

## Footnotes

1 Humanization in this context is the substitution of all or part of a rodent gene with the corresponding human gene and with inactivation, or knock out (KO), of the rodent gene. In our examples, substitution places a new human minigene under the control of the rodent promoter and simultaneously inactivates the rodent gene. Knock-out of the rodent gene and placement of the human gene in another genomic location, or under exogenous promoter control, may also be considered an example.

2 Active Genetic efficiency, AG%, is 100 times the frequency at which the parental genotype is transferred to the receiver chromosome. The receiver chromosome is the wild-type chromosome in a heterozygous configuration. Robust refers to efficiency that is high at all genetic loci to which the AG process is applied. Reusable means an AG process that can be used in successive generations without diminution in performance.

3 Active Genetic (AG) configuration – a configuration in an animal of Cas9, or another site-specific DNA hydrolase, and an AG cassette; the combination promotes the expression of Cas9 at a time in development wherein the chromosomal cut caused by Cas9 induces homology-directed repair and copying of a cargo gene from one chromosome to another.

4 Cas9 refers to any site-specific DNA hydrolase capable of expression in eukaryotic cells and includes all RNA-directed or non-RNA-directed site-specific DNA hydrolases, with or without homology to CRISPER-Cas9 genes, or any CRISPR-Cas9 gene from any organism.

5 AG cassette (AGC) – the cargo gene or marker of interest plus a guide-RNA (gRNA) cassette configured to promote expression of the gRNA. When the AGC is present in a cell also expressing Cas9, and preferably, but not exclusively, a cell undergoing meiosis prophase I, the Cas9-gRNA holoenzyme cuts the chromosome at a target site allowing integration and homology-directed repair (HDR) copying of the cargo human gene into the rodent target locus.

6 Our creation of the Del13 marker indel also created an identically located animal line, but with a Del24 marker indel. Due to time and animal number considerations, we used Del13 or Del24 configured FcgR1 loci interchangeably for many tests.

7 We also investigated the application of drug-regulated Cas9 systems to the AG problem. Both the DHRF and Tamoxifen regulated Cas9 system (here call ERT2-Cas9) (Liu et al., 2016; Maji et al. 2017) were investigated in preliminary lab testing in mouse and rat embryonic fibroblasts transfected cells treated with the regulator drug. DHFR-Cas9 constructs similarly arranged as Maji et al. exhibited trace levels of Cas9 and similar amounts in the presence or absence of drug; thus, we discontinue the investigation of this configuration. We used plasmid that encoded drug-regulated-ERT2-Cas9-P2A-GFP construct and transient transfection with a gRNA into rat embryonic fibroblasts using the ERT2 configuration Var30 of Liu et al., 2016 (Figure SM1). The ERT2-Cas9 showed a regulation ratio (+drug/-drug) of about 2500-fold, assessed using a Next-generation sequencing methodology (not shown). The ERT2-Cas9 drug-regulated system was selected for the development of rat and mouse strains to test its applicability as an accelerator strain necessary to promote the Active Genetics process. Similar rat and mouse versions of ERT2-Cas9 transgenic animals were prepared. The work on the rat strain was completed early in our efforts and characterization of the strain was conducted. Copious expression of ERT2-Cas9 was detected by immunohistochemistry in testicular cells of Cas9 heterozygous rats (not shown). However testicular cells from animals bearing an active genetic cassette did not exhibit Cas9 activity, using the T7 endonuclease assay conducted as shown in SR8, either before or after Tamoxifen was injected in the rats (not shown). Further tests using gRNA transfection testicular cells from the transgenic line were tested in cultures treated with 4-hydroxytamoxefin. In this test, using Next-Generation sequencing, we also did not observe Cas9 activity. Thus, biochemical Cas9 enzymatic was undetectable in ERT2-Cas9 bearing rats treated or untreated with tamoxifen. Based on these findings, further testing of Active Genetic activity in the ERT2-Cas9 rats and mice was stopped.

